# The STAT5-IRF4-BATF pathway drives heightened epigenetic remodeling in naïve CD4^+^ T cell responses of older adults

**DOI:** 10.1101/2021.08.27.457205

**Authors:** Huimin Zhang, Rohit R. Jadhav, Wenqiang Cao, Isabel N. Goronzy, Jun Jin, William J. Greenleaf, Cornelia M. Weyand, Jörg J. Goronzy

**Affiliations:** Department of Medicine, Stanford University, Palo Alto, CA 94305, USA; Department of Immunology, Mayo Clinic, Rochester, MN 55095, USA; Biochemistry and Molecular Biophysics, California Institute of Technology, Pasadena, CA 91125, USA; Department of Genetics, Stanford University, Palo Alto, Stanford, CA, USA; Department of Medicine, Division of Rheumatology, Mayo Clinic, Rochester, MN 55095, USA

**Keywords:** Naïve T cells, aging, immune memory, T cell activation, single cell multiomics, epigenomics, inflamm-aging

## Abstract

T cell aging is a complex process combining the emergence of cellular defects with activation of adaptive mechanisms. Generation of T cell memory is impaired, while a low-inflammatory state is induced, in part due to effector T cells. To determine whether age-associated changes in T cell fate decisions occur early after T cell activation, we profiled the longitudinal transcriptional and epigenetic landscape induced by TCR stimulation comparing naïve CD4^+^ T cells from young and older adults. In spite of attenuated TCR signaling, activation-induced remodeling of the epigenome increased with age, culminating in heightened BATF and BLIMP1 activity. Single cell studies, integrating ATAC-seq and RNA-seq data, identified increases in dysfunctional and in effector T cells and a decrease in BACH2-expressing memory cell precursors. STAT5 activation, in part due to a decline in HELIOS and aberrant IL-2 receptor expression, accounted for the induction of transcription factor networks favoring effector cell differentiation.

## Introduction

Aging is associated with a decline in immune function that manifests as an increased susceptibility to infection drastically highlighted in the recent pandemic of COVID-19 (Koff and Williams, 2020). Attenuated vaccination efficacy in the senior population against many pathogens, which aggravates the clinical consequences of infections (Gustafson et al., 2020), stresses the relevance of adaptive immunity for immune aging, in particular for CD4^+^ T cells, the major orchestrator of the response to most of the current vaccines. Simplified models, such as a decline in the total number of naïve T cells or contraction of TCR diversity due to reduced thymic activity have been found to be insufficient to explain age-associated immune dysfunction (Goronzy and Weyand, 2019). In fact, while the frequency of naïve CD4^+^ T cells declines with age and their TCR richness is reduced, the pool of naïve CD4^+^ T cells in human older adults is still large (Whiting et al., 2015) and diverse (Qi et al., 2014). In contrast to CD4^+^ T cells, the CD8^+^ compartment is more affected, with loss of naïve CD8^+^ T cells (Wertheimer et al., 2014), increased oligoclonal expansion and accumulation of virtual memory cells (Quinn et al., 2018) and specialized effector T cell population such as TEMRA (Pereira et al., 2020) and Taa cells (Mogilenko et al., 2021).

Immune aging constitutes a multidimensional process (Goronzy and Weyand, 2017; Pulendran and Davis, 2020). Given the non-linearity of a process like an immune response, the functional implications of a singular defect are difficult to predict. A prime example are the age-associated defects in TCR signaling that have been described in many studies (Fülöp et al., 1999, 2001; Li et al., 2012). Although recent RNA vaccines appear to overcome some of the defects in inducing an immune response (Anderson et al., 2020; Walsh et al., 2020), long-term memory is apparently not accomplished, reminiscent of earlier mouse studies that CD4^+^ T cell memory from young naïve cells functions well into old age, but memory cells generated from old naïve cells function poorly (Haynes et al., 2003). In vitro studies have shown that despite these stimulation defects, naïve CD4^+^ T cells from older human adults display a strong bias to differentiate into short-lived effector T cells although such a bias requires high signal intensity (Kim et al., 2018). Exogenous factors as well as epigenetic priming of the responding T cell population may contribute to this bias. It could explain the accumulation of effector T cell populations with age that contribute to the increased production of inflammatory mediators, coined as inflamm-aging.

System approaches integrating the different dimensions of functional pathways are needed to determine how age affects immune responses and to identify targets of interventions. Excellent work has been done in the murine system, including recent profiling of the aged immune system at single-cell resolution (Almanzar et al., 2020; Elyahu et al., 2019; Mogilenko et al., 2021). However, T cell homeostasis over lifetime is fundamentally different in mice and humans (den Braber et al., 2012), and age-associated changes in T cell population composition and cell function are therefore influenced by species differences (Zhang et al., 2021). Only one of these omics studies have examined whether the findings are relevant for human immune aging (Mogilenko et al., 2021). Moreover, static cross-sectional studies are only the first step and need to be followed by dynamic studies to identify the causality between age-associated changes. While such studies are more feasible in the mouse (Kaech et al., 2002; Kurachi et al., 2014), they may not always reflect the human system. To understand the complexity of the human CD4^+^ T cell response and to build a resource for subsequent studies, we monitored the trajectory of epigenetic and transcriptional changes of naïve CD4^+^ T cells from young and older adults after T cell receptor (TCR) stimulation, aiming to capture any aging signatures falling into the early kinetic window as candidates of rejuvenation targets.

By monitoring the epigenome and transcriptome of naïve CD4^+^ T cells upon TCR stimulation across a time course of 48 hours, we observed that T cells from older individuals had a higher propensity to TCR activation-associated chromatin changes despite having reduced TCR-induced phosphorylation of signaling molecules. Analysis of longitudinal changes identified STAT5 as the candidate priming transcription factor (TF) driving the excess TCR-induced differentiation exemplified by increased IRF4, BATF and BLIMP1 expression. Single cell multiome (scMultiome) profiling of activated naïve CD4^+^ T cells, integrating chromatin accessibility and transcriptomic data, revealed a subpopulation enriched with STAT5 activity that was increased in frequency with age. This early increased STAT5 activity was reflected by the upregulation of CD25 on naïve T cells at quiescent state, in part due to loss of the transcription factor HELIOS that represses *IL2RA* transcription. Inhibiting STAT5 signaling reoriented the epigenetic landscape of naïve CD4^+^ T cells in older individuals to that resembling young adults and restored the potential of memory formation by increasing TCF1 and reducing BLIMP1 expression.

## Results

### Effect of TCR signaling strength on epigenomic remodeling in naïve CD4^+^ T cells

Studies on the activation of human T cells have to rely on in vitro polyclonal stimuli that are generally supra-physiological and may therefore mask functionally important differences. Moreover, we wanted to control for differences in signaling strength as they occur with age. To mimic TCR signal inputs in the more physiological range, we stimulated T cells with polystyrene beads that had been coated with increasing amounts of anti-CD3 (αCD3) and a fixed amount of anti-CD28 (αCD28) Ab. The density of αCD3 on these beads was multifold lower than the commercial Dynabeads and induced different levels of signaling strengths. TCR stimulation was measured by phosphorylation of signaling molecules gating on the bead-bound cell population. Both, early signaling events as represented by ZAP70 and SLP76 phosphorylation (Figure S1A, left and middle panel) and downstream signaling events as represented by ERK phosphorylation (Figure S1C, right panel) correlated with the amount of αCD3 on the beads. αCD3 concentrations of 0.01 μg/mL that induced half-maximal ERK phosphorylation, and of 1 μg/mL, at which ERK phosphorylation just plateaued, were used in subsequent experiments as the low and medium TCR signal input, respectively. Of note, the TCR signal as measured by ERK phosphorylation peaked at 5 minutes and gradually tuned down over one hour (Figure S1B), which recapitulates the TCR signal induced by antigen-presenting cells (Yamane et al., 2005). The percentage of CD69^+^ CD4^+^ T cells at 24 hours of activation correlated with the initial TCR signal input (Figure S1C).

To examine epigenomic and transcriptional changes at different signaling strengths, we stimulated naïve CD4^+^ T cells from healthy individuals with αCD3-coated beads and performed ATAC-seq and RNA-seq at four longitudinal time points over 48 hours. Results at basal, low, and medium TCR signal input (0, 0.01 and 1 μg/mL αCD3) are shown as Uniform Manifold Approximation and Projection (UMAP) plots. The samples clustered according to the αCD3 concentration and post-activation time span (Figure 1A & 1B), demonstrating that the TCR stimulation-induced chromatin accessibility changes are dynamic and vary between different TCR stimulation strengths. Data from cells cultured with αCD28 Ab only did not show clear directionality in chromatin changes over time. Clustering at different time points after low-grade stimulation was not very succinct due to high inter-individual heterogeneity. Changes in chromatin accessibilities were minimal in the first 6 hours after stimulation; most changes occurred between 6 and 24 hours. In contrast, a major shift was seen in UMAP1 within one hour of higher intensity stimulation followed by a shift in UMAP2 over the next 24 to 48 hours (Figure 1A). Both, medium and low stimulation reached similar 48 hr endpoints at least in two individuals. The patterns were similar for transcriptome data (Figure 1B).

**Figure 1.**
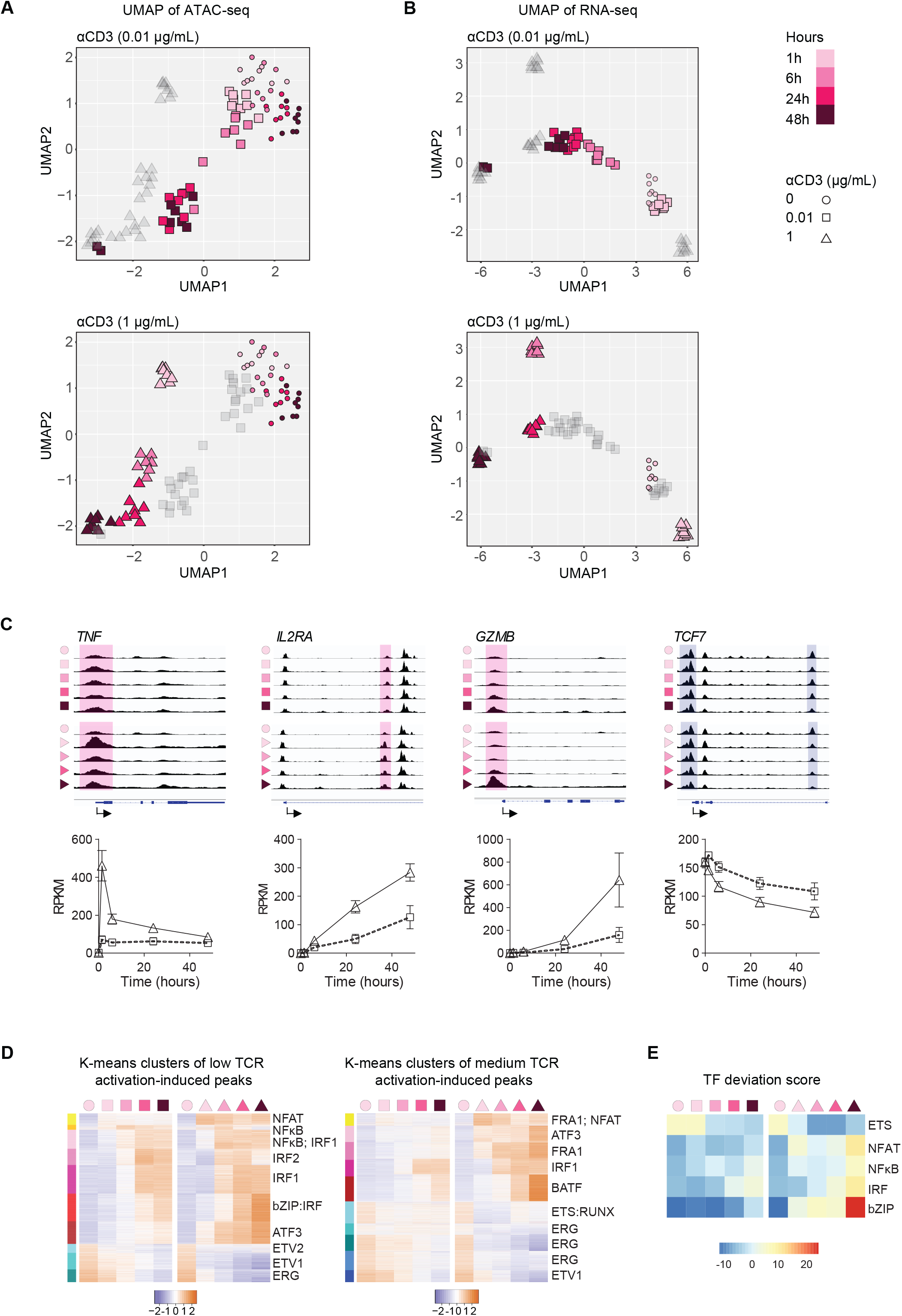
Chromatin accessibility changes induced by TCR stimulation of different signaling strengths. **A**. ATAC-seq UMAP visualization of naïve CD4^+^ T cells from eight healthy individuals stimulated with 0.01 μg/mL (squares) and 1 μg/mL αCD3-coated beads (triangles) for 1.5, 6, 24 or 48 hours. Time points are indicated by color code. Samples stimulated with 0 μg/mL αCD3 beads were included as control. **B**. RNA-seq UMAP visualization of the same set of samples. **C**. Aggregate genome tracks that close or open with TCR stimulation. Red shaded areas indicate peaks that open with TCR stimulation and blue areas peaks that close. Bottom panels are the corresponding gene expression data. **D**. Peaks changing in accessibility after low-(left) or higher-intensity TCR stimulation (right) were identified as described in Figure S2B. K-means clustering of the two peak sets are shown as heat plots: TFs with the highest binding motif enrichment in each cluster are indicated. **E**. ChromVAR deviation scores of selected TF motifs.

To gain insights on upstream regulators of the epigenomic changes, we identified differentially opened chromatin sites between TCR-stimulated T cells and those without stimulation (baseline) across the entire activation time course. The number of differentially open sites increased over time (Figure S2A). Cumulatively, about 5,000 sites changed in accessibility in response to low TCR signal and 30,000 sites to medium TCR signal (Figure S2B). Correspondingly, about 2000 genes were differentially expressed in response to the low and 6000 genes to the medium signal (Figure S2C & S2D). Typical activation-associated genes such as *TNF*, *IL2RA* and *GZMB* gained chromatin accessibility after TCR stimulation and quiescence-associated genes such as *TCF7* lost accessibility (Figure 1C). Longitudinal transcript expression closely trailed changes in chromatin accessibility. K-means analysis of differentially open sites at low TCR signal input identified seven clusters that gradually opened after TCR stimulation (yellow/red-labeled clusters) and three that gradually closed (cyan/blue-labeled clusters) (Figure 1D, left heat plot). Sites identified to change with low stimulation changed in a similar pattern after high intensity stimulation, with the major difference that accessibility changes occurred earlier in the time course. Transcription factor (TF) motif enrichment analysis suggested that the opening clusters are mainly regulated by the NFκB, NFAT, IRF and bZIP families while those closing by the ETS family. At low TCR input, the NFAT and NFκB-induced chromatin opening was apparent as early as after one hour of TCR activation, followed by sites enriched for IRF and bZIP motifs (Figure 1D, left heat plot). This time course is consistent with the previously reported fast activation of NFAT upon T cell-APC interaction (Marangoni et al., 2013).

Many additional sites changed accessibility at higher intensity stimulation (Figure 1D, right heat plot). These sites can be grouped into five clusters with increasing and five with decreasing accessibility. The dominant TF motifs gaining in accessibility were those of the bZIP family. NFAT, together with ATF/bZIP-induced sites preceded that of IRF and BATF/bZIP. Enrichment for IRF motifs, while present in one cluster, was less dominant than for low-intensity stimulation. All the T cell activation-associated TFs show a clear dichotomy of ChromVAR deviation between low and medium TCR signal input (Figure 1E). Taken together, the differential TCR signal inputs induced dynamic and signal strength-dependent chromatin accessibility changes in naïve CD4^+^ T cells.

### T cells from older adults are more poised to chromatin accessibility changes after TCR stimulation

TCR signaling has been shown to be blunted with age, in part due to a reduced expression of miR-181a and the associated increased concentrations of several phosphatases (Li et al., 2012). Consistent with these studies, we found reduced ERK and LAT phosphorylation in naïve CD4^+^ T cells from older individuals stimulated with 1 μg/mL αCD3-coated beads (Figure 2A). We therefore expected activation-induced epigenomic changes in T cells from older individuals to be shifted to those seen with lower signaling strength in young adults. Our study population shown in Figure 1 included four 21-35 year-old adults and four adults older than 65 years (Supplement table 1). To identify age-dependent shifts in the activation-induced epigenetic state of naïve CD4^+^ T cells, we performed principal component analysis (PCA) of the 5000 most variable sites and plotted the result separately for samples from young and older individuals stimulated with different TCR signals across the 48-hour time course. Results are shown for PC1, which explained 50% of the variance (Figure 2B), while other PCs accounted for less than 8% of variance each. For cells cultured with beads lacking αCD3 Ab, both young and older individuals had minimal longitudinal changes along the time course. At low TCR input, all individuals showed an increase in PC1 starting from 6 hours of stimulation while at medium TCR input, the increase started as early as at 1.5 hours, documenting that T cell activation strength correlated with epigenetic changes as shown by PC1 scores (Figure 2B). Notably, older individuals maintained at least an equal PC1 score at both TCR inputs at all time points compared with those from young adults. This is in contrast to the blunted TCR signaling, suggesting that the activation-induced chromatin state and early signaling events are uncoupled with age.

**Figure 2.**
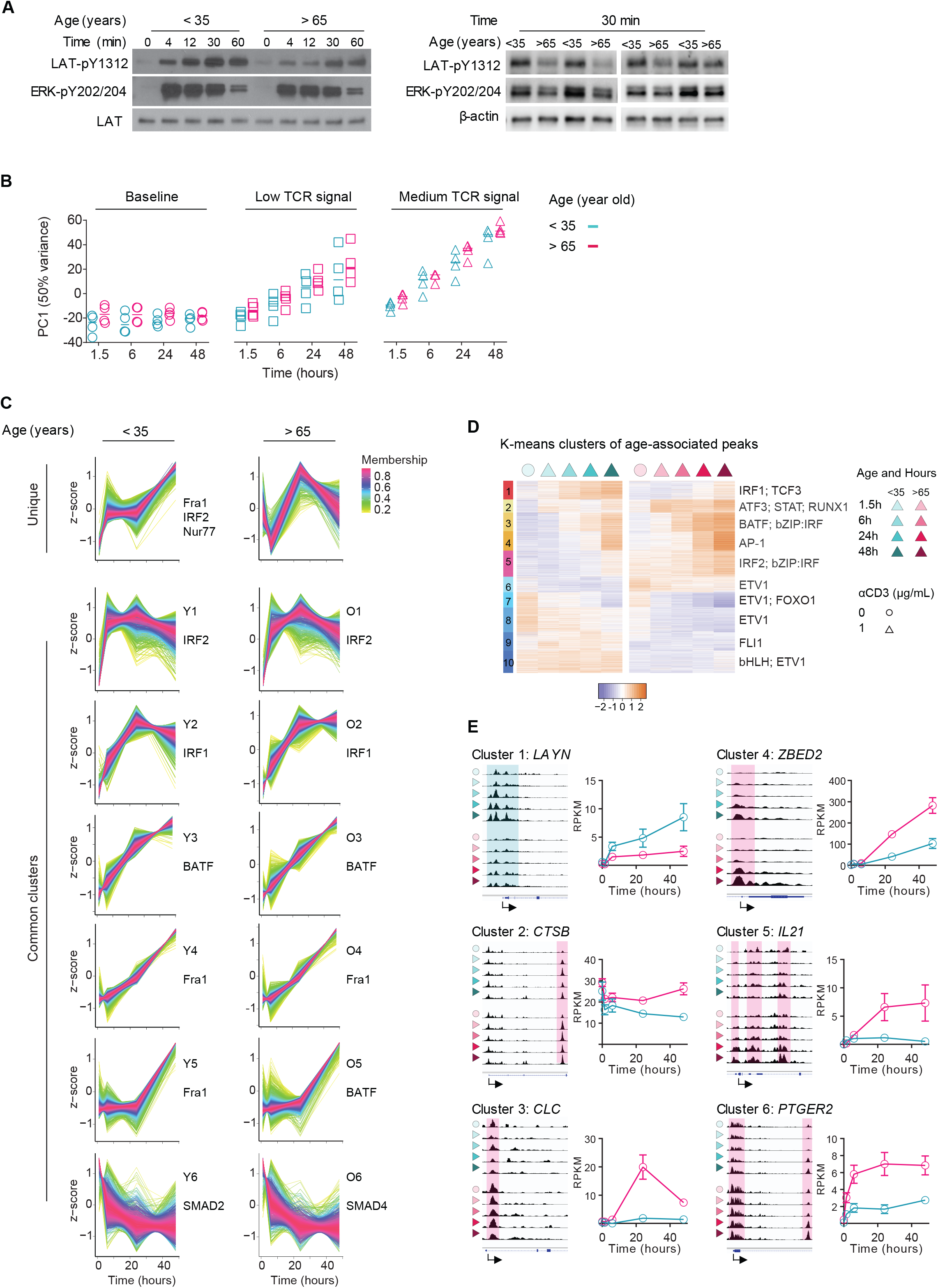
Dissociation of TCR-induced signaling and chromatin accessibility changes in naïve CD4^+^ T cells from older adults. **A**. LAT and ERK phosphorylation was measured from 0 to 60 minutes after stimulation of naïve CD4^+^ T cells with polystyrene beads coated with 1 μg/mL αCD3 (left panel). Phosphorylation at 30 minutes was measured for 4 young and 4 older individuals (right panel). **B**. PCA was performed on the 5000 most variable sites of the ATAC-seq dataset described in Figure 1A. PC1, accounting for 50% variance is shown separately for stimulated naïve CD4^+^ T cells from young and older adults. **C**. Peak sets from ATAC-seq data of young and older adults were generated that significantly differed from baseline at any of the post-stimulation time points. Peaks from younger and older adults were separately subjected to TCseq clustering. Results are shown for seven clusters with different temporal pattern, six of them grossly similar between young and old. Membership scores are indicated by color code. TF motif enrichment was determined by HOMER for each cluster on sites with membership scores of >0.6. TFs with the highest binding motif enrichment are indicated, detailed results are given in Figure S4A. **D**. K-means clustering of peaks differential between young and older individuals at each time point after higher intensity TCR stimulation (see Figure S3). TFs with the highest binding motif enrichment for each cluster are indicated. **E**. Aggregate genome tracks of representative sites of cluster 1-6 in D (left panel). Magenta shaded areas indicate peaks that open more in older adults and cyan ones peaks that are open more in young adults. Gene expression from RNA-seq data (right panel).

Cumulatively over the time course after activation with higher intensity stimulation, 3078 sites had higher and 2557 lower accessibility in T cells from older compared to young adults. To a large extent these peaks included sites that were differentially accessible at lower stimulation (Figure S3), indicating that the higher intensity stimulation was more informative to examine age-related differences. In our subsequent analysis, we therefore focused on this dataset.

Since TCR-induced chromatin changes were longitudinally orchestrated (Figure 1D), we sought to first determine whether the response in older individuals had unique kinetic features. We used fuzzy k-means clustering on all sites changing with stimulation to identify temporal patterns of activation-induced chromatin changes in each age group (Wu and Gu, 2021). Results for seven clusters (from a range of 2 to 10) were chosen as the minimal number capturing different patterns. In Figure 2C, clusters are ordered by the appearance of early changes in the time course. Six clusters were largely similar between the two age groups, with only small kinetic differences, suggesting that the progression of chromatin changes after T cell activation was not grossly influenced by age. Clustering correlated with enrichment for selective TFs; the most enriched TF are shown in Figure 2C and a more detailed analysis in Figure S4A. A cluster unique for young adults was characterized by an early rapid change followed by a plateau for 24 hours before further differentiation. Biological pathway analysis showed that the annotated genes of this cluster were enriched for the T cell activation pathway (Figure S4B), and TF motif enrichment analysis suggested regulation by FRA1 (which has a similar motif as AP1), NF-κB and NUR77 (Figure S4A), all consistent with the increased phosphorylation of TCR signaling molecules in young adult. The two next clusters (clusters 1 and 2) in the time course with changes plateauing at 6 and 24 hours, respectively were enriched for IRF family member motifs. Sites in clusters 3-5 were progressively opening over the entire time course. Corresponding genes were enriched for developmental pathways (Figure S4B). Motif analysis suggested that bZIP family members are the dominant upstream regulators. Transcript analysis supported the regulation of temporal patterns by different TFs as implicated by motif enrichment (Figure S4C). *NR4A1* (encoding NUR77) peaked early. *IRF1* and *IRF4* peaked slightly later, followed by *IRF9* with slightly different patterns in young and old. bZIP family TFs came in several waves. Fra1 (encoded by FOSL1) and AP1 (encoded by *FOS* and *JUN*) was rapidly upregulated in the first hour and *ATF5* and *BATF* were progressively upregulated throughout the 48-hour time course (Figure S4C). Interestingly, the clusters with more sustained activation progression after 24 hours all showed higher enrichment for bZIP family motifs in T cells from older adults (Figure 2C & S4A), consistent with an elevated expression of *BATF* in older adults (Figure S4C). A similar pattern was observed for BLIMP1 encoded by *PRDM1*. Sites of cluster 6, closing immediately after TCR stimulation were regulated by SMAD family members and enriched for T cell activation and adhesion pathways. The unique cluster in old adults did not show significant enrichment of any TF motif (Figure S4A).

To directly focus on age-associated signatures, we performed k-means clustering of the differential peaks between young and older individuals at each time point. Gap statistics suggested to cluster into ten groups including 5 with sites gradually opening with TCR stimulation and 5 clusters that closed (Figure 2D). Most of the closing sites were already distinct at baseline and regulated by members of the ETS family, suggesting these aging signatures are imprinted in the quiescent state. Increased accessibility in older individuals was activation-induced and sites were enriched for STAT, bZIP and bZIP:IRF family motifs. Only one cluster with sites opening after activation was less accessible in older adults. This cluster was regulated by the TCF and IRF families. TCF is important for T memory cell development, suggesting that the response pattern of T cells in older adults to TCR activation includes compromised memory formation. Accessibility tracks of genes representative for each cluster are shown together with the corresponding transcript data (Figures 2E and S5A). Accessibility differences at each of the time points correlated with the fold difference in transcripts at 48 hours (Figure S5B).

### Accelerated chromatin changes in T cells from older adults are driven by the STAT5 pathway

The propensity of T cells from older adults to undergo accelerated TCR-induced chromation changes cannot be explained by TCR signaling that is blunted (Figure 2A); only chromatin changes in the first hours after TCR stimulation appeared to be more prominent in the young (Figure 2C). To identify alternative potential upstream regulators that precede the increased accessibility to bZIP family motifs and that may poise the larger response in older adults, we identified enriched TF binding motifs in the whole peak set using ChromVAR and plotted TFs with deviation score differences in the top 10 percentile separately for young and older individuals (Figure 3A). The TF clustering patterns resembled that of differentially opened sites, falling into two categories: one with TF binding sites closing across the time course, e.g. the ETS family; and one with TF binding sites opening, e.g. the STAT, IRF, bZIP families. Of note, TFs driving opening sites trended to show higher deviation scores in older individuals, corroborating the notion that naive CD4^+^ T cells from older adults are more prone to increasing chromatin accessibilities after TCR stimulation (Figure 3A). Specifically, STAT5 motif enrichment plateaued as early as at 6 hours, while that of IRF4 at 48 hours and that of BATF was still rising at 48 hours (Figure 3B). These data suggest that STAT5 may be the prime TF that drives elevated activation-induced chromatin changes in older adults.

**Figure 3.**
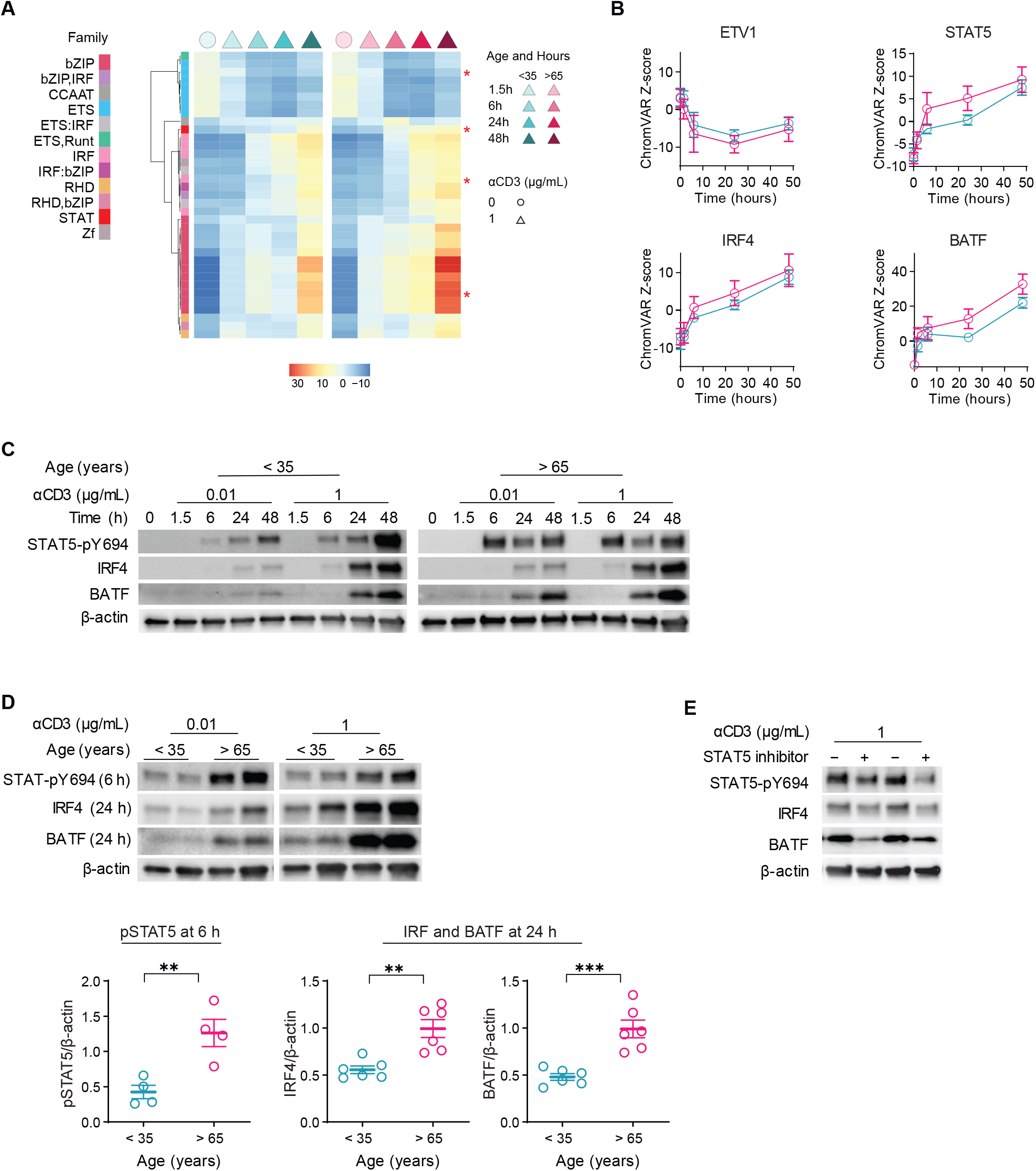
Accelerated chromatin changes in T cells from older adults are driven by the STAT5 pathway. **A**. Heatmap representation of top 35 most variable TFs based on ChromVAR deviation scores of the ATAC-seq data as described in Figure 1A. Each column is the score median of the four young (left) and the four older (right) individuals. Stars indicate the representative TFs in B. **B**. ChromVAR deviation scores for representative TFs from Figure 3A are shown as mean+SEM. **C**. Phosphorylated STAT5 (pSTAT5), IRF4 and BATF were measured at indicated times after TCR stimulation. Western blots are shown for one pair of one young and one older individual representative of three experiments. **D**. Representative blots of pSTAT5 measured at 6 hours and IRF and BATF at 24 hours after low and higher intensity stimulation (top). Summary results for 4 to 6 young and older adults after median intensity stimulation shown as mean±SEM (bottom). Data were analyzed with unpaired t-test. **p < 0.01, ***p < 0.001. **E**. Naïve CD4^+^ T cells from older adults were activated for 48 hours with indicated αCD3 beads in the presence of DMSO or a STAT5 inhibitor. Western blots for pSTAT5, IRF4 and BATF are from 2 experiments representative of 5.

To confirm the kinetic sequence of increased TF activities in older adults, we examined the protein level of implicated TFs. Consistent with the temporal pattern in ChromVAR, phosphorylated STAT5 (pSTAT5) emerged at 6 hours followed by IRF4 and BATF in both young and older individuals (Figure 3C). In particular, pSTAT5 peaked already at 6 hours in the old whereas it displayed minimal activity in young adults. In CD4^+^ T cell responses from older individuals, pSTAT5 was significantly higher after 6 hours (p<0.01), while IRF4 and BATF only after 24 hours of stimulation (p<0.01, Figures 3D). Blocking STAT5 phosphorylation by a STAT5 inhibitor reduced upregulation of IRF4 and BATF expression (Figure 3E), corroborating that STAT5 drives the accelerated chromatin changes in naïve CD4^+^ T cells from older adults.

### A subpopulation of naïve CD4^+^ T cells with increased STAT5 activity is expanded with age

Multiple lines of evidence suggest that the naïve T cell compartment is not a homogeneous cell population (Davenport et al., 2020). The increased epigenetic TCR response may therefore reflect a distinct subset that increases with age. We employed scMultiomics to investigate population heterogeneity of resting and activated naïve CD4^+^ T cells at the epigenome and transcriptome level. The cluster of 18-hour TCR-stimulated naïve CD4^+^ T cells was distinctly separate from resting cells in ATAC-seq UMAP and confirmed by *CD69* expression (Figure S6). Within activated cells, four subclusters could be distinguished, most obviously, when ATAC-seq and RNA-seq data were integrated (Figure 4A). TF-binding motif analysis revealed that clusters 1 was enriched for KLF4, cluster 2 for STAT5 and clusters 3 and 4 for BACH2- and NFAT-binding sites (Figure 4B, top panel). TF-binding footprints confirmed increased STAT5 activity in cluster 2, Bach2 in cluster 3 and 4 and NFAT activity in cluster 4. Enrichment for activation-induced gene expression differed dependent on the cluster. Cluster 1 was characterized by high transcript numbers of *CXCR4*, which was rapidly downregulated after T cell activation in our bulk RNA-seq data. Pathway analysis shows cluster 1 was enriched in senescence-associated genes and relatively resistant to T cell activation. Thus, we defined cluster 1 as low responders. Cluster 2 showed high transcript numbers of *PRDM1* and was enriched for RUNX3-regulated immune responses and thus was defined as effector precursors. Cluster 3 was enriched for TCF1 and BACH2 targets, both of which are involved in the differentiation of memory precursors. Cluster 4 showed high expression of *CD69*, *TNF* and *IL2*, all of which are signs of rapid T cell activation, and thus was defined as fast responders (Figure 4B).

**Figure 4.**
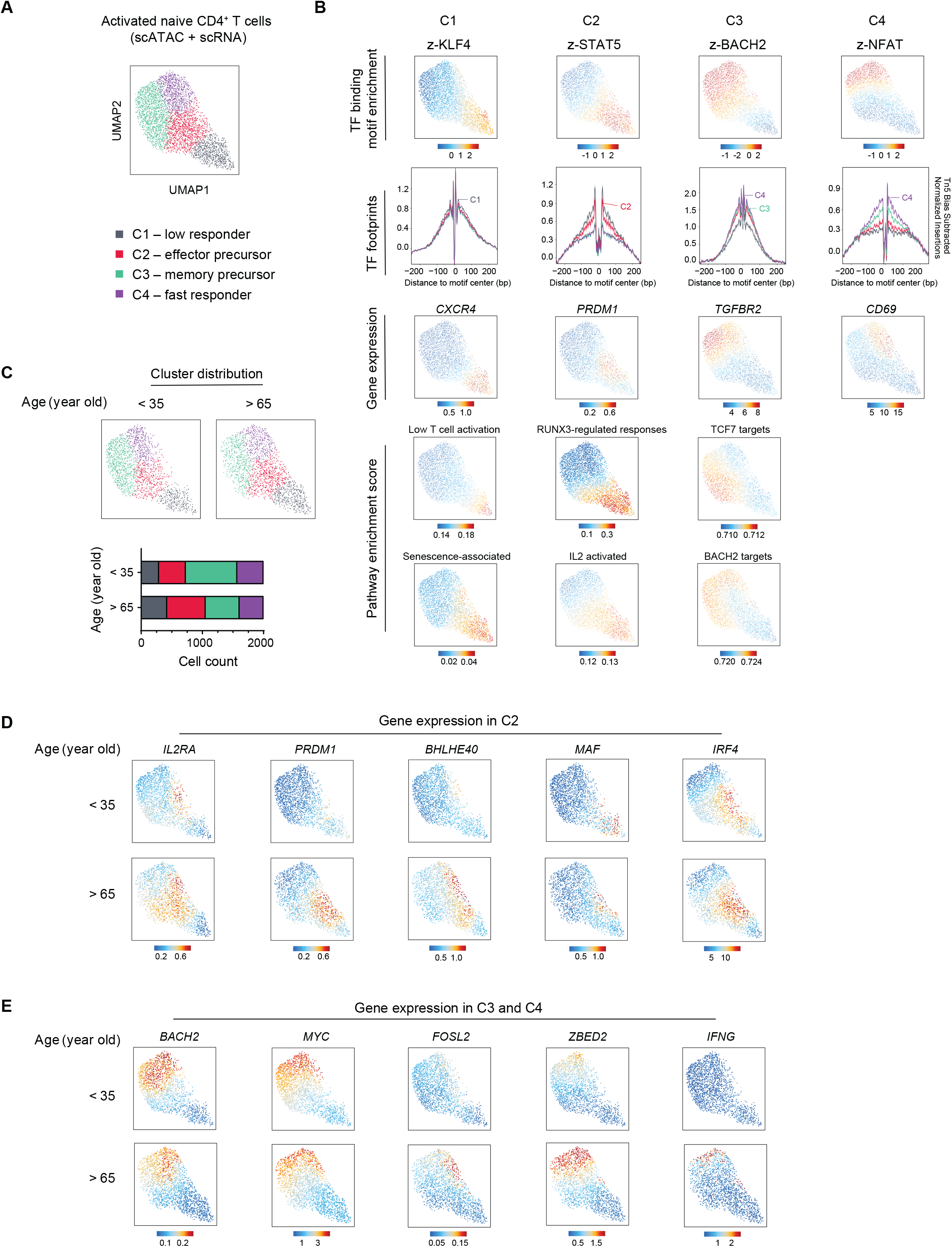
Age-associated expansion of a naïve CD4^+^ T cell subset with increased activation of the STAT5 pathway. **A**. Naïve CD4^+^ T cells from two young and two older individuals were left unstimulated or activated with beads coated with 1 μg/mL αCD3 and αCD28 for 18 hours. Nuclei were subjected to combined scATAC/scRNA-seq. Activated cells were identified in UMAP maps and confirmed by CD69 expression (see Figure S6A). UMAP of integrated scATAC-seq and scRNA-seq data are shown with each point representing one cell. Clusters indicated by different colors were generated using Surat graph clustering approach. **B**. Representative TFs in each cluster are shown as ChromVAR deviation scores projected on the UMAP map and as footprints using the color code for the different clusters in Figure 4A (top two panels). Representative gene expression and indicated pathway enrichment score in each cluster are projected on the UMAP (bottom two panels). **C**. scMultiomic UMAPs of activated naïve CD4^+^ T cells from young and older adults are presented separately (top panel). Frequencies of cells for different clusters are shown as stacked bar graphs (bottom panel). **D**. Imputed gene expression in Cluster 2 with differential expression for young and older adults. **E**. Genes in cluster 3 and 4 with differential expression for young and older adults.

To explore whether the excess in activation-induced chromatin modifications in older adults was representative of the whole naïve CD4^+^ T cell compartment or reflected distinct subpopulations, we plotted the UMAPs of activated single cells of young and older individuals separately (Figure 4C). Older individuals had notably larger effector precursor and low-responder clusters, both enriched for STAT5 binding sites (Figure 4B & 4C). Moreover, cells from older adults in cluster 2 were characterized by increased expression of CD25 and of several transcription factors involved in T effector cell differentiation (*IRF4*, *PRDM1*, *BHLHE40*, and *MAF*) (Figure 4D). The upregulation of CD25 was already detectable in unstimulated cells (Figure S7B & S7C), suggesting that the age-associated elevation of STAT5 signaling is imprinted in the quiescent state. However, age-associated differences were not limited to an expansion of cells and a corresponding increased gene expression in cluster 2. The memory precursor cluster, enriched for BACH2 binding sites, was smaller in older individuals (Figure 4B & 4C) and *BACH2* expression was reduced in activated CD4^+^ T cells from older adults (Figure 4E). *BACH2* has been implicated in maintaining T cell quiescence and favoring memory over effector cell development (Roychoudhuri et al., 2016; Tsukumo et al., 2013; Yao et al., 2021). The lower expression of *BACH2* in older adults could therefore render T cells from older adults more susceptible to activation and may contribute to the accelerated accessibility to bZIP motifs as shown in Figure 2D. Similarly, expression of *MYC* was largely limited to clusters 3 and 4 and was increased in T cells from young adults suggesting greater metabolic changes. Conversely, clusters 3 and 4 had increased expression of the TF *ZBED2* in T cells of older adults. ZBED2 has been shown to antagonize IRF1 function (Somerville et al., 2020), which may negatively affect TH1 differentiation (Kano et al., 2007) and contribute to T cell dysfunction (Li et al., 2019). Of note, only very few cells, all in cluster 4, had evidence of epigenetic or transcriptional hallmarks of differentiated memory cells, such as of the *IFNG* gene, confirming that the expansion of cluster 2 or the accelerated chromatin changes in the older adults do not reflect expanded memory cells phenotypically masquerading as naïve cells (Pulko et al., 2016). Taken together, scMultiomic data and protein expression data collectively revealed a subpopulation of naïve CD4^+^ T cells with increased STAT5 signaling that accumulated with age and accounted for the higher propensity to TCR stimulation-associated epigenetic changes.

### CD25 upregulation is induced by loss of *IKZF2* with age

Since CD25 was upregulated with age at both transcript and protein level (Figure S7B & S7C), we examined whether the chromatin accessibility at the *IL2RA* gene locus was altered with age. Comparing ATAC-seq of unstimulated naïve CD4^+^ T cells from young and older adults, we observed two enhancer regions more open in naïve CD4^+^ T cells from older adults (Figure 5A). One of the enhancers (enhancer A) has been reported to be associated with autoimmunity risk by influencing the timing of gene responses to extracellular signals (Simeonov et al., 2017). We transfected the reporter luciferase construct containing enhancer A into naïve CD4^+^ T cells from young and older adults and found it to induce 30% higher luciferase activity in older adults (Figure 5B). Enhancers are occupied by transcription factors (TFs) for gene activity control (Spitz and Furlong, 2012). We identified TFs binding *IL2RA* enhancers from the UCSC genome browser (Figure S8A) (Kent et al., 2002; Rosenbloom et al., 2013). Some of the identified TFs are activated in response to extracellular stimuli such as FOS and BHLHE40, while others are constitutively active in the resting state, including TCF1, IKZF1 and IKZF2 (ENCODE Project Consortium, 2012). Since we observed age-associated increased accessibility at *IL2RA* in the resting state and IKZF2 is the resting state TFs exhibiting the most enrichment at both *IL2RA* enhancer regions (Figure 5A & S8A), we examined the occupancy of HELIOS (encoded by *IKZF2*) at both enhancers by ChIP-qPCR. HELIOS showed a two-fold increase in binding in naïve CD4^+^ T cells from young compared to older adults (Figure 5C), indicating it may confer repression of *IL2RA* promoter activity. We transfected the reporter construct containing the *IL2RA* enhancer A together with an *IKZF2* cDNA construct to force overexpression of HELIOS and found reduced promoter activity in the presence of HELIOS (Figure 5D). This was further corroborated by a negative correlation between *IKZF2* and *IL2RA* transcripts in naïve CD4^+^ T cells sorted for CD25 expression (Figure S8B). When comparing transcript and protein levels in naïve CD4^+^ T cells from young and older individuals, HELIOS was found to be significantly diminished with age (Figure 5E & 5F). Moreover, silencing *IKZF2* in resting naïve CD4^+^ T cells from young adults upregulated CD25, reproducing the patterns seen in old adults (Figure 5G). Taken together, we found *IKZF2* transcription was lost with age in naïve CD4^+^ T cells resulting in CD25 upregulation. Since *IKZF2* is traditionally recognized as a TF maintaining the stability of regulatory T cells (Kim et al., 2015; Thornton et al., 2010), these data also confirm that the subset of CD25^lo^CD4^+^CD45RA^+^ T cells that is increased with age and in part accounts for the enhanced epigenetic changes are not regulatory T cells.

**Figure 5.**
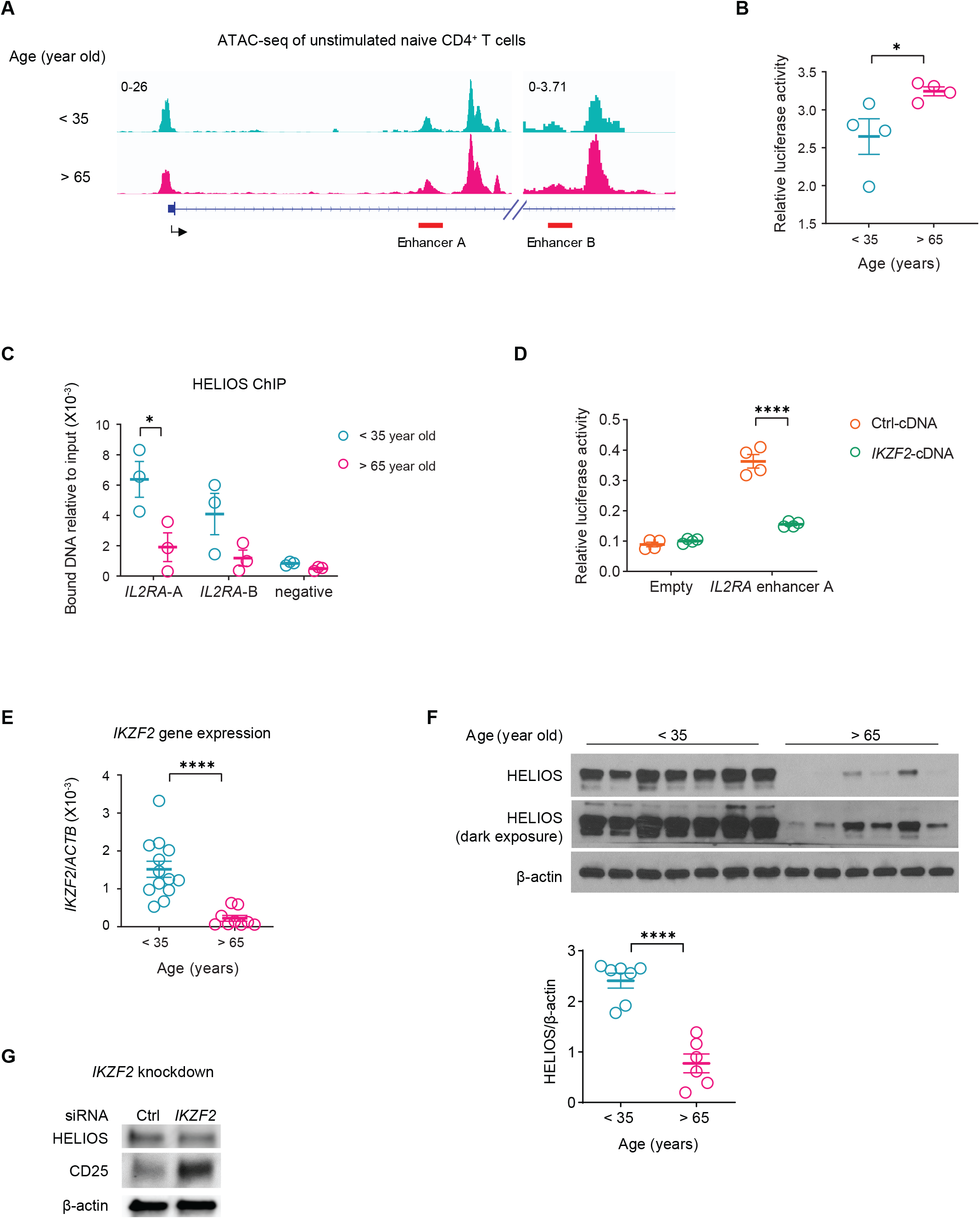
CD25 upregulation is induced by a loss of *IKZF2* with age. **A**. Aggregate genome tracks of unstimulated naïve CD4^+^ T cells from 4 young and 4 older individuals at *IL2RA* enhancer locus (GRCh37, Chr10:6,091,304-6,111,593 and Chr10:6,085,160-6,090,232). **B**. Luciferase activity in unstimulated naïve CD4^+^ T cells from 4 young and 4 older individuals after transfection of a reporter construct containing *IL2RA* enhancer A. **C**. ChIP-qPCR with HELIOS (encoded by *IKZF2*) - specific antibody of unstimulated naïve CD4^+^ T cells from 3 young and 3 older adults. **D**. Luciferase activity in unstimulated naïve CD4^+^ T cells after transfection of reporter construct with or without *IL2RA* enhancer A together with control cDNA or *IKZF2* cDNA. **E**. *IKZF2* gene expression in unstimulated naïve CD4^+^ T cells from 13 young and 9 older adults. Data were analyzed with unpaired t-test. ****p < 0.0001. **F**. *IKZF2* protein expression in unstimulated naïve CD4^+^ T cells from 7 young and 6 older adults. Data were analyzed with unpaired t-test. ****p < 0.0001. **G**. CD25 expression in unstimulated naïve CD4^+^ T cells transfected with control or *IKZF2* siRNA.

### Inhibition of IL2-STAT5 signaling rewires naïve CD4^+^ T cells from older adults to a younger epigenetic state

Since 40% of naïve CD4^+^ T cells in older individuals (vs 20% in young) had upregulated CD25 (Figure S7C), we examined whether early STAT5 activity after TCR stimulation resulted in the excess epigenetic changes. We used either IL-2 receptor (IL-2R) blocking antibodies or a STAT5 inhibitor in a 48 hour stimulation of naïve CD4^+^ T cells from older individuals and assessed the epigenetic changes by ATAC-seq. PCA of sites significantly affected by the interventions showed that both IL-2R blocking and STAT5 inhibition shifted the chromatin accessibility states as defined by PC1 in the direction towards those of younger individuals (Figure 6A). TF motif enrichment analysis of sites driving PC1 yielded STAT and bZIP binding motifs (Figure 6B). 25% of differentially opened sites induced by IL-2R blocking or STAT5 inhibition were also differentially open between young and old adults (indicated by dark red, Figure 6C & S9A); examples included opening peaks in *TCF7* and *BACH2* and closing peaks in *IRF4* and *BHLHE40* (Figure 6D & S9B). Consistent with STAT5 ChIP-seq data of human naïve CD4^+^ T cells (Liao et al., 2011), we found STAT5 binding at the *IRF4* and *BHLHE40* genes by ChIP-qPCR in naïve CD4^+^ T cells from older adults after 48-hour TCR stimulation (Figure 6E). Inhibition of IL2-STAT5 activity downregulated expression of IRF4 and BHLHE40 at the protein level (Figure 6F). To determine whether the increased STAT5 activity accounted for the preferential effector over memory CD4^+^ T cell generation previously reported for T cell responses of older adults (Jin et al., 2021; Kim et al., 2018), we determined TCF1 (encoded by *TCF7*) and BLIMP1 (encoded by *PRDM1*) expression after 5 days of TCR activation. pSTAT5 inhibition in cultures with naïve CD4^+^ T cells from older adults downregulated BLIMP1 and upregulated TCF1 (Figure 6G), recapitulating the relationship between young and older individuals (Figure 6H). Taken together, by dampening IL2-STAT5 signaling, naïve CD4^+^ T cells from older adults can be rewired to exhibit a chromatin accessibility state after TCR activation that resemble that in young adults with increased potential of memory formation.

**Figure 6.**
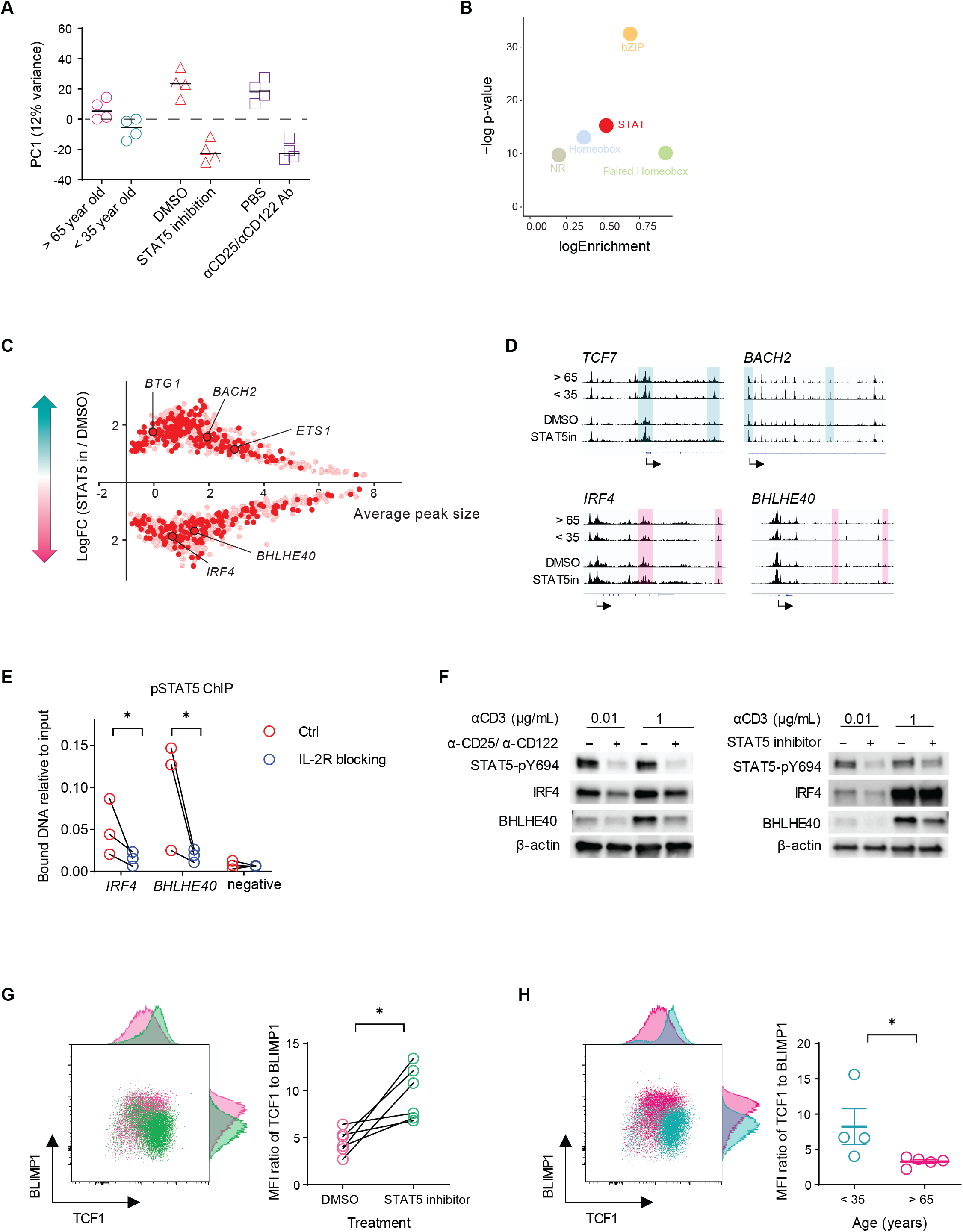
Inhibition of IL2-STAT5 signaling rewires activated naïve CD4^+^ T cells from older individuals to a younger epigenetic response pattern. **A**. Four older individuals were activated in the presence of either DMSO control, STAT5 inhibitor, PBS control or IL-2R-blocking antibodies for 48 hours and then subjected to ATAC-seq. Data of the four treatments were subjected to PCA together with the 48 hours samples from the four young and four older individuals without any treatment described in Figures 1 and 2. Results for PC1 are shown. **B**. TF binding motif enrichment of PC1 loading sites in A. **C**. LogFC of differential peaks between STAT5 inhibition and DMSO treatment (p<0.05) are plotted against average peak size. Peaks that are also differentially accessible between young and older individuals are indicated in dark red. **D**. Accessibility tracks of representative genes. Magenta-shaded areas indicate peaks that are more open in older adults or with DMSO treatment; cyan indicates peaks more open in young adults or with STAT5 inhibition treatment. **E**. ChIP-qPCR with pSTAT5 antibody of 48-hour activated naïve CD4^+^ T cells treated with either PBS or IL-2R blocking antibodies. Data were analyzed with paired t-test. *p < 0.05. **F**. pSTAT5, IRF4 and BHLHE40 were measured by Western blot in 48-hour activated naïve CD4^+^ T cells for each treatment conditions. **G**. Naïve CD4^+^ T cells from 6 older adults were activated with 1 μg/mL αCD3-coated beads for five days together with DMSO or STAT5 inhibitor. TCF1 and BLIMP1 expression was measured by flow cytometry. The MFI ratio of TCF1 to BLIMP1 was plotted. Data were analyzed with paired t-test. *p < 0.05. **H**. Naïve CD4^+^ T cells from 4 young and 5 older adults were activated for five days. TCF1 and BLIMP1 protein expression was assayed as in Figure 6G. The MFI ratio of TCF1 to BLIMP1 is plotted. Data were analyzed with unpaired t-test. *p < 0.05.

## Discussion

Here, we describe that, in spite of attenuated proximal TCR signaling, naïve CD4^+^ T cells from older adults respond to activation with accelerated remodeling of the epigenome, culminating in increased BATF and BLIMP1 activity. This epigenetic response pattern explains why generation of long-lived memory T cells is impaired with age and activated naïve T cells are biased to differentiate into effector T cells that may be short-lived and contribute to mediators driving inflamm-aging. Single cell studies integrating ATAC-seq and RNA-seq revealed a subset of activated T cells with increased STAT5 activity in older adults, in part due to the aberrant expression of CD25. The increased STAT5 activity accounted for the induction of TF networks that were driving the increased chromatin accessibility in older adults. Importantly, addition of a STAT5 inhibitor prevented this epigenetic response pattern and induced TCF1 instead of BLIMP1, a constellation favoring the generation of long-lived memory and TFH cells. These findings underscore the potential of single cell combined ATAC-seq and RNA-seq studies to unravel pathways in complex processes and to identify targets for interventions.

The advent of multiomic technologies has enabled the exploration of the human immune systems at unprecedented scale and resolution. Most of the human studies have been cross-sectional focusing on constitutively expressed marker profiles, thereby resulting in a more detailed description of cell types such as B cell subsets (Glass et al., 2020) or the classification of human dendritic cells (Villani et al., 2017). In the context of aging, longitudinal studies of the transcriptome, proteome and metabolome in young and old individuals over a period of two years have identified profiles predictive for different aging trajectories (Ahadi et al., 2020). Multiomic studies of biological process responses have the potential of providing mechanistic insights and targets for interventions but are challenging in humans. Investigators have tried to overcome limitations by integrating results from murine and human data, such as when characterizing T cell development trajectories (Chopp et al., 2020; Le et al., 2020). Species-specific differences such as those in T cell homeostasis limit this approach for better defining T cell aging. We have therefore focused here on a system of in vitro activation of human naïve CD4^+^ T cells to define the temporal patterns of epigenetic and transcriptional changes after activation and define regulatory networks that cause age-associated differences. We were careful to adapt the in vitro activation system to more physiological ranges of TCR signaling strength and to avoid supraphysiological stimulation. In these studies, the integration of ATAC-seq and RNA-seq data, in particular at the single nuclei level, was particularly insightful and allowed conclusions on regulatory networks.

Based on the widely described, attenuated phosphorylation of TCR signaling molecules in T cells from older adults that we also confirmed in our activation system, we expected epigenomic changes to be reduced by age and predicted that their induction require increased stimulation strength. In contrast to our expectations, activation-induced remodeling of chromatin accessibility was increased with age and age-associated differences culminated in the increased expression and activity of BATF, across a range of signaling strength. In the LCMV model, BATF activity occurs early after infection, mostly in conjunction with IRF4, and has been implicated in the differentiation and survival of early CD8^+^ effector T cells (Kurachi et al., 2014). A recent scRNA-seq study tracking the differentiation trajectory of LCMV-specific CD8^+^ T cells suggested a divergence of effector and memory signatures at the first cell division event (Kakaradov et al., 2017). We focused on a time window up to 48 hours, a time point when naïve CD4^+^ T cells after TCR stimulation just started dividing. We observed that the increased BATF activity in older T cells is not the first age-related event but preceded by pSTAT5 and IRF4 signals in the first 6 to 24 hours after activation. Also already at these early time points, PRDM1 expression (encoding BLIMP1) was increased that is known to direct effector cell differentiation at the expense of memory and TFH cell generation (Johnston et al., 2009; Rutishauser et al., 2009). Only one temporal pattern of chromatin accessibility changes was more prevalent in T cell responses of young adults. This pattern involved rapid changes in the first one to six hours after activation involving sites enriched for bZIP family (presumably AP1) and NUR77 motifs, likely reflecting early events due to superior TCR signaling in the young.

The TF networks implicated to drive the differential chromatin accessibility changes in T cells from older adults provide an explanation for earlier functional studies that naïve CD4^+^ T cells from older adults preferentially develop into effector T cells rather than memory or TFH cells. The increased pSTAT5 activity accounted for the induction of TF networks and was at least in part due to increased expression of the IL-2 receptor. Heightened IL-2 signaling is known to favor terminal effector differentiation in CD8^+^ T cells (Pipkin et al., 2010), thereby compromising memory cell formation (Kalia et al., 2010; Pipkin et al., 2010). In CD4^+^ T cells from older adults, this bias towards effector cell differentiation was mechanistically linked to increased expression of miR-21 repressing the expression of several negative regulators including PTEN, SPRTY and PDCD4 (Kim et al., 2018). Besides direct regulation of effector cytokines, STAT5 also induced miR-21 in malignant T cells (Lindahl et al., 2016). Moreover, STAT5 inhibits the differentiation of TFH cells via the BLIMP-1-dependent repression of BCL6 (Johnston et al., 2012). In addition to impairing TFH development, the reduced BCL6 activity derepresses the ecto-ATPase CD39 that is expressed on effector and particularly exhausted T cells (Cao et al., 2020). Increased expression of CD39 is a hallmark of T cell responses of older individuals, thereby modifying purinergic signaling and causing adenosine-mediated immunosuppression and cell death (Fang et al., 2016).

Integrated scRNA- and scATAC-seq analysis showed increased pSTAT5 activity in two clusters of cells that were both increased in cell numbers with age. Both clusters also shared the enrichment for senescence-like pathways. Cluster 2 cells had heightened CD25 expression, consonant with previous observations in unstimulated total T cells of older adults (Pekalski et al., 2013). The mechanisms leading to increased pSTAT5 activity in Cluster 1 are unclear, however, IL-2 receptor inhibition prevented accelerated chromatin remodeling in CD4^+^ T cell responses of older adults as did STAT5 inhibition. Cluster 2 cells had several features of effector cells, including PRDM1 expression, RUNX3 pathway activity and presence of CD39^+^ cells. Importantly, they do not include fully differentiated effector cells as recently described (Pulko et al., 2016). *IFNG* transcript-expressing cells were very few, exclusively assigned to Cluster 4 of rapidly responding cells. Beyond these differences related to STAT5 activity, scATAC/RNA-seq also showed an age-associated decrease in BACH2 and increase in ZBED2 expression in cluster 3 and 4 cells. BACH2 has been shown to suppress effector memory-related genes and enforce the transcriptional and epigenetic programs of stem-like CD8^+^ T cells, at least in part by controlling access of AP-1 factors to enhancers (Roychoudhuri et al., 2016; Tsukumo et al., 2013; Yao et al., 2021). Thus, BACH2 deficiency may be an additional mechanism to select against the generation of long-lived memory cells. ZBED2 were found enriched in dysfunctional CD8^+^ T cells (Li et al., 2019) and elevated *ZBED2* expression in older adults may contribute to the compromised long-term memory.

Excess STAT5 activity in activated naïve CD4^+^ T cells can contribute to the inflammatory milieu in older adults through several layers of regulation. STAT5 activity in CD4^+^ T cells has been implicated in promoting autoimmune disorders including experimental autoimmune encephalomyelitis (Sheng et al., 2014) and airway inflammation (Fu et al., 2020) through upregulation of effector cytokines, such as GM-CSF and IL-9. Consistently, we observed transcript levels of both cytokines increased with age. Consonant with these observations are our previous findings that older adults have a bias towards TH9 cell generation through upregulated IRF4 and BATF (Hu et al., 2019). Increased effector T cell differentiation driven by IL2 may therefore be an important regulator of the low-grade inflammatory state in older adults, also coined as inflamm-aging. Identification of IL2 as an upstream regulator of age-related epigenetic differences in T cell responses revives the discussion on the relative benefits and harm of this inflammatory state. Inflammation can reinvigorate exhausted T cells (West et al., 2013) and sustained STAT5 activity can promote anti-tumor activity through both CD4^+^ and CD8^+^ T cells (Ding et al., 2020; Zhou et al., 2021).

## Acknowledgements

This work was supported by the National Institutes of Health (R01 AR042527, R01 HL117913, R01 AI108906, R01 HL142068, and P01 HL129941 to C.M.W, and R01 AI108891, R01 AG045779, U19 AI057266, and R01 AI129191 to J.J.G). The content is solely the responsibility of the authors and does not necessarily represent the official views of the National Institutes of Health. We thank Dr. Peng Li from the National Institutes of Health for providing the processed ChIP-seq data file of human CD4^+^ T cells. We thank Dr. Fabian Müller for suggestions on single cell data analysis. We thank the Stanford Genome Sequencing Service Center and Novogen for providing sequencing services.

## Conflict of Interest Statement

The authors declare that the research was conducted in the absence of any commercial or financial relationship that could be construed as a potential conflict of interest.

## Author Contributions

H.Z., R.R.J., W.J.G., C.M.W. and J.J.G. designed research and interpreted data. H.Z., W.C. and J.J. performed experimental work. R.R.J. and I.N.G. analyzed high-throughput data. H.Z., R.R.J. and J.J.G. wrote the manuscript.

## Data Availability Statement

Raw sequencing data have been deposited in SRA with the BioProject accession # PRJNA757466.

## STAR Methods

### Study design

For sequencing studies, 16 individuals were recruited who did not have an acute or active chronic disease or a history of cancer or autoimmune diseases. Chronic diseases such as hypertension were permitted if controlled by medication. The studies were approved by the Stanford University and Mayo Clinic Institutional Review Boards, and all participants gave informed written consent. Basic demographic information is listed in Supplement table 1. Naïve CD4^+^ T cells from subjects 1-8 were stimulated for four time points with polystyrene beads coated with 0, 0.01 and 1 μg/mL αCD3. All 96 samples were collected for ATAC-seq. RNA-seq was performed on the 8 baseline samples and the 64 post-stimulation samples. Naïve CD4^+^ T cells from subjects 9-12 were stimulated with polystyrene beads (αCD3=1 μg/mL) for 18 h and collected for scMultiome sequencing. Naïve CD4^+^ T cells from subjects 13-16 were stimulated with polystyrene beads (αCD3=1 μg/mL) for 48 h in presence of either PBS, 5 μg/mL of αCD25 and αCD122 Ab, DMSO or 50 μM of STAT5 inhibitor (Cayman chemical, #15784) in DMSO. For experiments not involving genome-wide sequencing, buffy coats from de-identified donors (n=105, age 21 to 35 years or over 65 years were purchased from the Stanford Blood Bank.

### Naïve CD4^+^ T cell isolation and activation

PBMC were Ficoll-isolated from buffy coats. Naïve CD4^+^ T cells were isolated by negative selection with EasySep™ Human Naïve CD4^+^ T Cell Isolation Kit (Stemcell, #19555). For time course study, cells were rested overnight in RPMI medium (Sigma, # R8758) and then stimulated with polystyrene beads labeled with αCD3 at 0, 0.01 or 1 μg/mL in 96 well round bottom plate. Each well contained 200,000 cells and an equal number of beads in 200 μL RPMI medium. TCR stimulation started with a centrifugation of the plate for 3 min at 500 g. At indicated time, cells were washed with PBS and subjected to ATAC-seq, RNA-seq or scMultiome experiments.

### Polystyrene beads labeling

Streptavidin-coated polystyrene beads (Bangs laboratories, #CP01006) were washed with 1XPBS/1%FBS and mixed with biotinylated αCD28 (Miltenyi Biotec, #130-093-386) at 1 μg/mL and biotinylated αCD3 (BioLegend, #317320) at 0, 0.01 or 1 μg/mL in 10X PBS/1%FBS for 20 min. Labeled beads were washed twice with 1XPBS/1%FBS and resuspended in RPMI medium.

### IKZF2 knockdown in unstimulated naïve CD4^+^ T cells

Cells were transfected with either ON-TARGETplus siRNA negative control (Horizon, #D-001810-10-20) or siRNA targeting IKZF2 (Horizon, #L-006946-00-0005) using the Amaxa Nucleofector system and P3 Primary Cell 4D-Nucleofector™ X Kit (Lonza, # V4XP-3024). 2 hours after transfection, cells were collected and resuspended in prewarmed medium and cultured for 2 days prior to harvest and analysis.

### Flow cytometry

Surface protein labeling was performed by mixing cells with antibodies in 1XPBS/1%FBS. For phosphorylated protein staining, cells were fixed (BD, #554655), permeablized (BD, #558050), and stained with antibodies. For TF staining, cells were fixed (BD, #554655), permeablized (eBioscience, #00-8333-56), and stained with antibodies (Supplement table 2). Stained samples were acquired with BD LSR Fortessa and analyzed with FlowJo v10.

### ATAC-seq

50,000 cells were collected for standard transposition reactions (Buenrostro et al., 2015) and sequenced with NovaSeq 6000 by Genome Sequencing Service Center in Stanford University. The ATACseq reads were preprocessed to remove adapters followed by filtering for read quality cutoff of 20. The reads were further filtered to select only autosomal reads for further analysis. The reads were then mapped to the hg19 genome using bowtie. Peak calling was performed using macs followed by generating a consensus peak set present in at least 3 samples with more than 50% site overlap. Reads were then assigned to the consensus peaks using feature counts and used for downstream analysis.

Samples were separated into groups based on the αCD3 concentration, timepoint and age. Principle component analysis (PCA) and uniform manifold approximation and projection (UMAP) was performed on counts normalized using variance stabilizing transformation (vst) followed by removing batch effect for donors (except for aging comparison) from DESeq2. Deviations in transcription factor accessibility patterns were estimated on all consensus peaks using ChromVar (Schep et al., 2017). Differential accessibility was assessed on data quantile normalized using voomWithQualityWeights from limma with addition of offsets from conditional quantile normalization (CQN) (Hansen et al., 2012) to account for GC content bias. We used duplicateCorrelation to control for donor effects followed by fitting a robust linear regression model with sample correlation blocking for donors. The differences between sample groups were estimated by fitting contrasts to the model followed by empirical Bayes moderation. We used contrasts to identify aggregate differences across all time points or age groups while controlling for baseline changes at each timepoint. The identified differential sites were further median normalized across sample groups and used to identify clusters using k-means clustering. Gap statistic was used to determine number of clusters in the data with a cutoff at 10 clusters. The differential sites were annotated with associated genes using GREAT and used for downstream analysis and comparisons with RNAseq (McLean et al., 2010). Gene ontology analysis was done using ChIPseeker (Yu et al., 2015).

### RNA-seq

150,000 cells were collected for RNA extraction. RNA libraries were generated with universal plus mRNA-seq kit (Nugene, #0520-A01) and sequenced with NovaSeq 6000 by Novogene. The RNAseq reads were preprocessed using the nf-core rnaseq pipeline v1.4.2 (Ewels et al., 2019) to determine reads mapped to genes in GRCh37 genome. The preprocessing for downstream analysis was kept consistent with ATACseq as follows. For PCA and UMAP, the RNAseq read counts were also normalized by “vst” followed by removing batch effect for donors (except for aging comparison) using DESeq2. To identify differential transcripts, we used CQN offsets along with quantile normalized data from voomWithQualityWeights from limma. We also used duplicate Correlation to control for donor effects followed by fitting a robust linear regression, blocking for donor and accounting for sample correlation. The differentially expressed genes between groups were estimated by fitting contrasts followed by empirical Bayes moderation. The contrasts were setup to identify aggregate differences in transcripts controlling for only baseline changes. The differential genes were further median normalized and used for k-means clustering. Gap statistic was used to determine number of clusters in the data with a cutoff of 10 clusters.

### TCseq

Fuzzy k-means cluster analysis of the ATAC-seq dataset was conducted using the TCSeq (Time Course Sequencing Data Analysis) package in R (Wu and Gu, 2021). First, the set of sites for analysis was defined as the union of peak locations detected by MACS2 from all time points. This set was then filtered for genomic regions that showed a significant change in read counts over time. Differential read count analysis used a negative binomial generalized linear model implemented by edgeR. Finally, the time course of normalized read counts at each genomic interval were clustered using soft clustering (’fuzzy k-means’). To reduce the bias introduced by difference in absolute values, the data was standardized using a z-score transformation prior to clustering. We tested a range of starting seed clusters (2-10) to find the optimum number of clusters for capturing distinctive temporal patterns in the data.

### Single cell Multiome (scATAC- and scRNA-seq)

Naïve CD4^+^ T cells were left unstimulated or stimulated for 18 hours with 1 μg/mL αCD3-coated beads. 40,000 cells were collected and subjected to low input nuclei isolation. A total of 10,000 nuclei from subject 9 and 10 were pooled for library generation and 10,000 nuclei from subject 11 and 12 (Supplement table 1). scATAC and gene expression libraries from each pool were generated at the Stanford Genomics Core following the Chromium Next GEM Single Cell Multiome ATAC + Gene Expression User Guide and sequenced with NovaSeq 6000 by Novogene. The multiome data for scRNAseq and scATAC seq was processed using Cellranger ARC followed by ArchR (Granja et al., 2021). Pathway enrichment score were calculated by Ucell package in R using the following datasets from MSigDB (Liberzon et al., 2011; Subramanian et al., 2005): low T cell activation (GSE22886) (Abbas et al., 2005), RUNX3-regulated genes (Reactome: R-HSA-8949275), TCF7 targets (Kolmykov et al., 2021), Senescence-associated genes (Reactome: R-HSA-2559584), IL-2 activated genes (GSE8685) (Marzec et al., 2008), BACH2 targets (Kolmykov et al., 2021).

### ChIP-qPCR

Chromatin immunoprecipitation (ChIP) assays were performed using the ChIP-IT kit (53040, Active Motif) according to the manufacture’s instruction. Briefly, 5 × 10^6^ cells were fixed with fixation buffer (containing 1% formaldehyde), washed and sonicated on ice by sonicator (Active Motif) to obtain 100-1000 bp DNA fragments. Fragmented DNA was immunoprecipitated with pSTAT5 and HELIOS Ab (Supplement table 2). Immunoprecipitated DNA were purified and subjected to qPCR using the following primers:

IRF4 (1-AGTTGCAGGTTGACCTACGG, 2-TTCGATCGTCTGAGATGCTG);
BHLHE40 (1-GCCTGTTGACACAACGTCAC, 2-GATCAGGTTTCTGCTGACGC);
IL2RA-A (1-TGATCCGTATCTTGCCTTCC, 2-GAAACTCCAGGGCAACAAAG);
IL2RA-B (1-CCACCCACTCTTTGCTGGAT, 2-TCAATGGGTAACAGCACCAGT);
Negative control (1-AACCTGCAAAACATGGTTATTT, 2-AATTTGCCCAAACAGCAAGT).

### qPCR

RNA was extracted with RNeasy Micro Kit (Qiagen, #74004) and reverse-transcribed with High-Capacity cDNA Reverse Transcription Kit (ThermoFisher, #4368814). Quantitative PCR (qPCR) was performed with PowerUp™ SYBR™ Green Master Mix (ThermoFisher, #A25776) and the following primers:

IKZF2 (1-GCTCCTCGCTGAAGATGGAG, 2-TGCCTAACGTGTGTTTGTGC);
IL2RA (1-ATTTCGTGGTGGGGCAGATG, 2-TCTCTTCACCTGGAAACTGACTG).

### Luciferase reporter assay

HEK-293T cells were cotransfected with empty pGL4.23 (Promega, # E8411) or pGL4.23-IL2RA CaRE4 plasmid (Simeonov et al., 2017), and Renilla luciferase reporter plasmid, a HELlOS-expressing plasmid (Sino Biological, #HG19573-UT), or a control vector (Sino Biological, # CV011). After 48 hours, luciferase activities were measured by the Dual-Luciferase Reporter Assay System (Promega, # E1910) according to the manufacturer’s instructions. For measuring IL2RA enhancer activity in human T cells, pGL4.23-IL2RA CaRE4 and Renilla plasmid were cotransfected into unstimulated naïve CD4^+^ T cells using the Amaxa Nucleofector system and P3 Primary Cell 4D-Nucleofector™ X Kit (Lonza, # V4XP-3024), cells then were cultured with IL-7 for 3 days, and subjected to luciferase activity measurement.

### Western blot

Cells were lysed with RIPA buffer (ThermoFisher, #89900) supplemented with protease and phosphatase inhibitors (ThermoFisher, #78441). Protein lysates were loaded on 4-15% precast TGX gels (BioRad, #4561086) and transferred to nitrocellulose membranes (BioRad, #1704270). Membranes were blocked with 5% milk in PBST buffer and incubated at 4 °C overnight with primary antibodies (Supplement table 2). After washing twice with PBST buffer, membranes were incubated with secondary antibodies (Cell Signaling, #7074) for 1 h and washed twice. Chemiluminescent signals were developed with SuperSignal™ West Femto Maximum Sensitivity Substrate (ThermoFisher, #34095).

## Notes

### Competing Interest Statement

The authors have declared no competing interest.

