## Supplement Figures and Tables for "The STAT5-IRF4-BATF pathway drives heightened epigenetic remodeling in naïve CD4^+^ T cell responses of older adults"

**Figure S1. Titration of TCR stimulation, related to Figure 1.**

**A.** Naïve CD4+ T cells were stimulated with polystyrene beads coated with indicated amounts of αCD3 and 1 μg/mL of αCD28 for 5 minutes. Mean fluorescence intensity (MFI) of phosphorylated ZAP70, SLP76 and ERK was measured by flow cytometry gating on bead-bound cells. **B.** MFI of ERK phosphorylation at 1 μg/mL of αCD3 was measured from 0 to 60 minutes. **C.** Frequency of CD69+ cells were determined by flow cytometry after 16 hours of TCR stimulation with indicated αCD3 concentrations. Experiments shown are representative of four experiments.

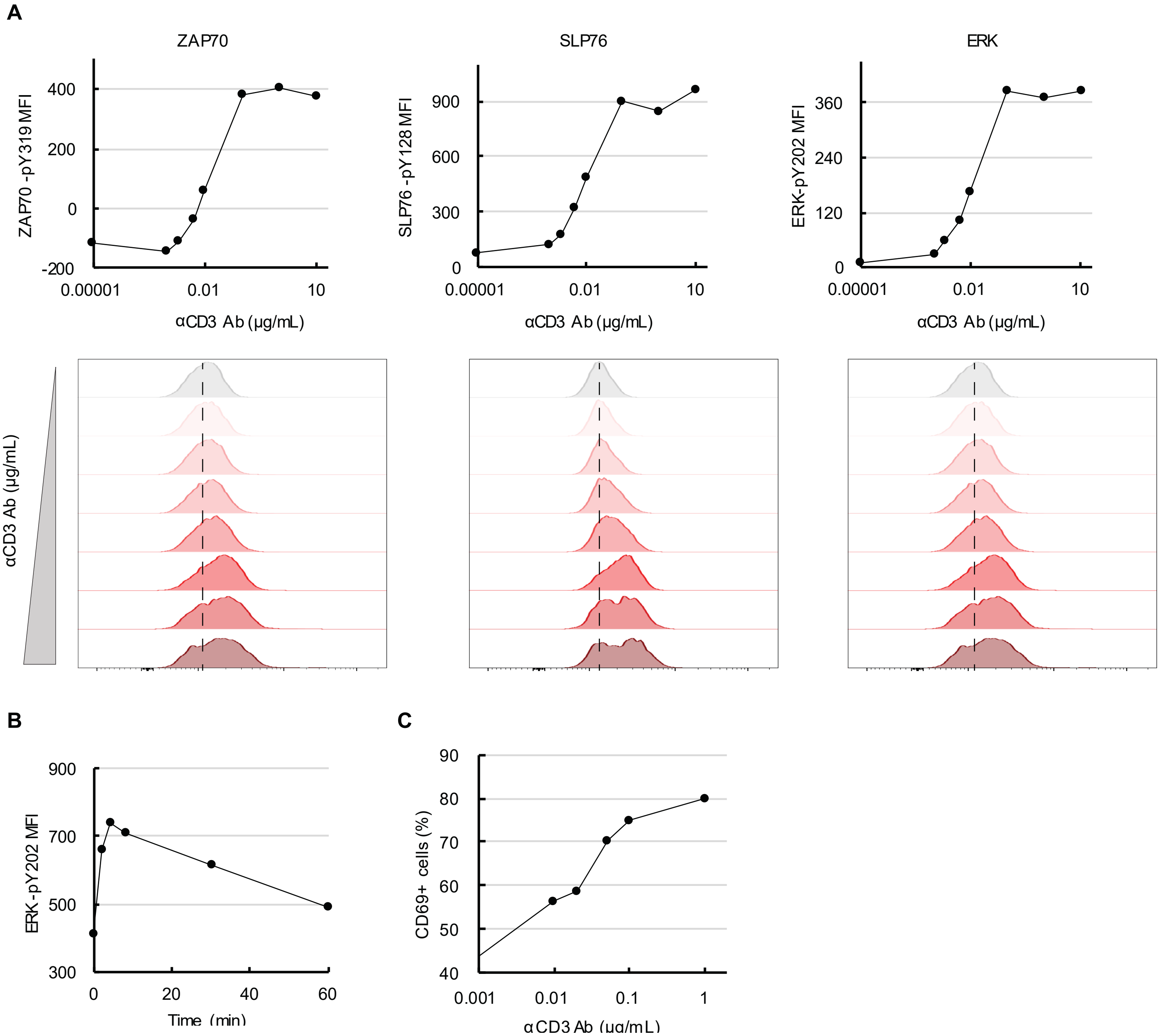

**Figure S2. Longitudinal epigenomic and transcriptomic changes induced by TCR stimulation**

Naïve $CD4^+$ T cells from eight individuals were stimulated with 0.01 (low, square) or 1 μg/mL (median, triangle) αCD3-coated beads and subjected to ATAC-seq and RNA-seq at indicated time points. **A.** Bar graphs show the number of peaks more or less accessible in activated cells compared to base line at each time point. **B.** Total differential peaks between low (left) or high (right) TCR stimulation and baseline are plotted as average $\log_2$ fold-change (logFC) versus $\log_2$ mean reads per peak. Peaks with p-values below 0.05 are indicated in red. **C.** Bar graphs show the number of significantly up- or down-regulated gene transcripts for each time point ($p < 0.05$). **D.** Total up- or down-regulated gene transcripts in activated cells compared to baseline plotted as logFC versus the average expression.

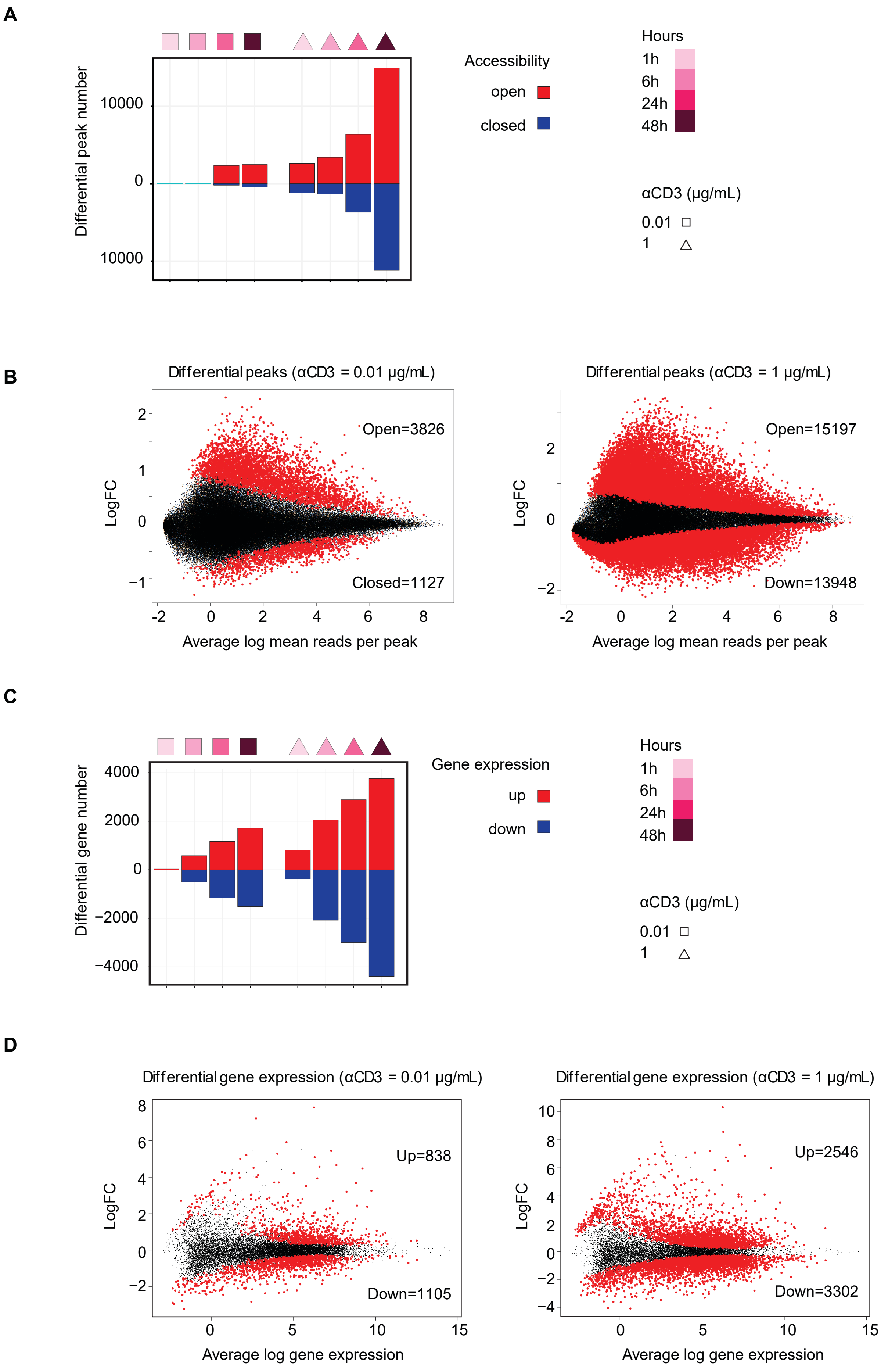
A
Differential peak number
10000
0
10000
Accessibility
open
closed
Hours
1h
6h
24h
48h
αCD3 (µg/mL)
0.01
1
B
Differential peaks (αCD3 = 0.01 µg/mL)
Open=3826
Closed=1127
LogFC
Average log mean reads per peak
Differential peaks (αCD3 = 1 µg/mL)
Open=15197
Down=13948
LogFC
Average log mean reads per peak
C
Differential gene number
4000
2000
0
−2000
−4000
Gene expression
up
down
Hours
1h
6h
24h
48h
αCD3 (µg/mL)
0.01
1
D
Differential gene expression (αCD3 = 0.01 µg/mL)
Up=838
Down=1105
LogFC
Average log gene expression
Differential gene expression (αCD3 = 1 µg/mL)
Up=2546
Down=3302
LogFC
Average log gene expression

**Figure S3. Higher intensity TCR stimulation amplifies age-associated chromatin accessibility differences seen at baseline or with low intensity stimulation**

Sites more accessible ($p < 0.05$) in naïve $CD4^+$ T cells from young (left) or old (right) adults were identified at baseline and at several time points after TCR stimulation with beads coated with low and medium concentrations of αCD3. Relationships between the three peak sets are shown as Venn diagrams.

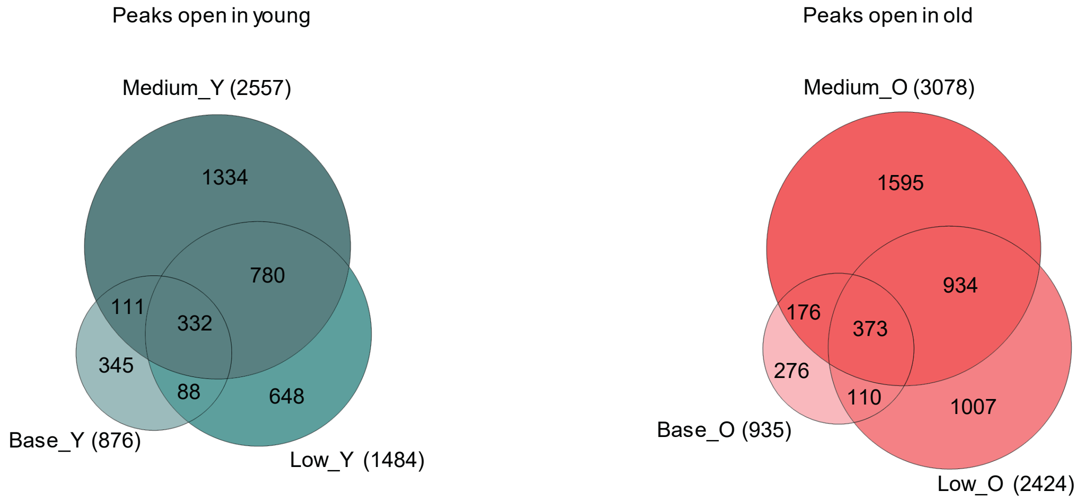

**Figure S4. related to Figure 2C. Activation-induced temporal patterns of epigenetic changes in naïve CD4$^+$ T cells from young and older adults.**

**A.** Peak sets in clusters with different temporal patterns and membership scores >0.6 as shown in Figure 2C were analyzed for TF motif enrichments by HOMER. Graphs show the fold enrichment (X-axis) in each cluster from young (to the left) and old (to the right) adults plotted vs the significance level (Y-axis). Five TFs with the lowest p-values are shown. **B.** Genes assigned to differentially accessible regulatory sites were examined for significant enrichment in biological pathways. Selected GO terms associated with the different temporal patterns in young (upper panel) and older adults (lower panel) are shown. **C.** Gene expression time course of indicated TFs from RNA-seq data.

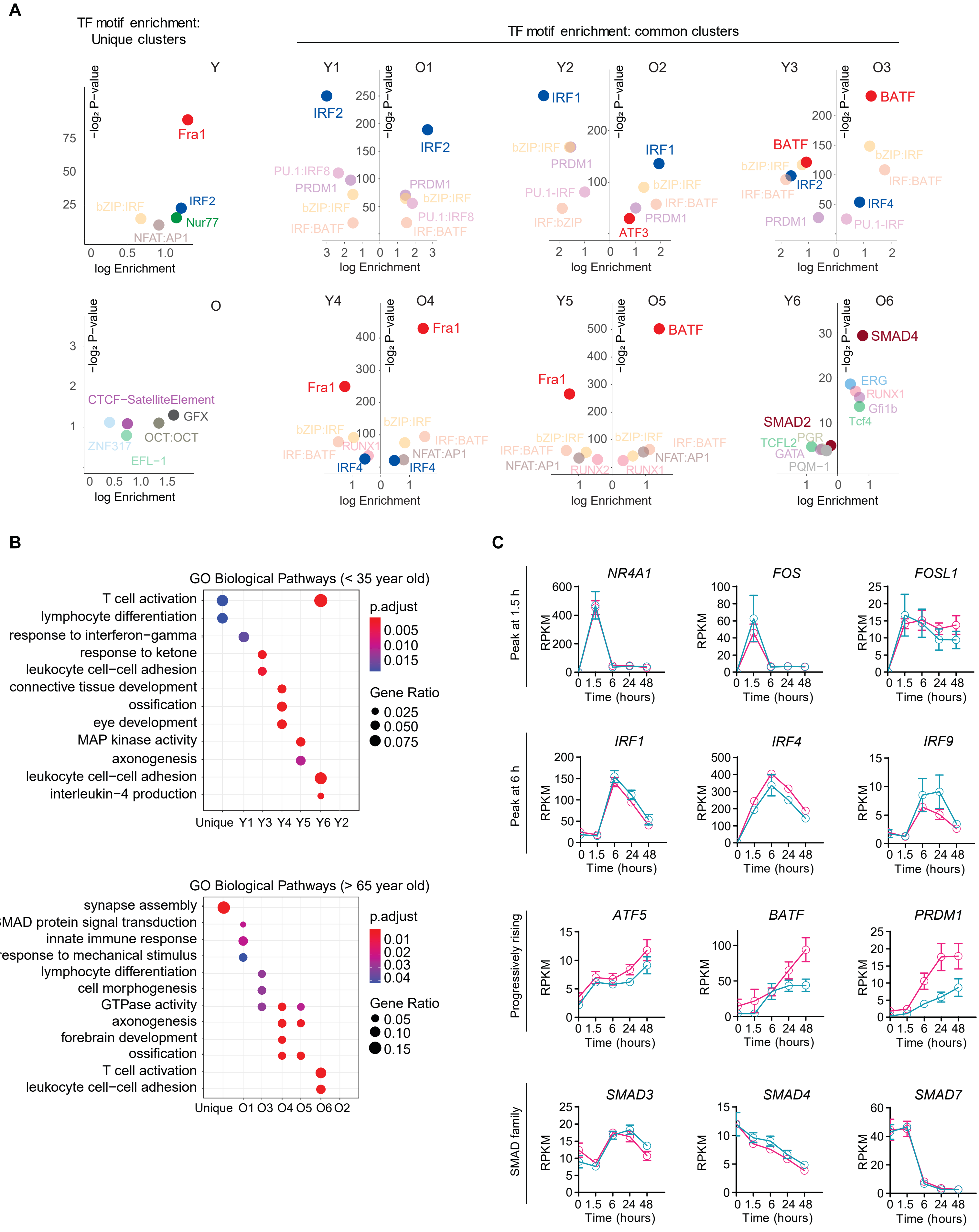
A
TF motif enrichment: Unique clusters
TF motif enrichment: common clusters
Y
Fra1
bZIP:IRF
IRF2
Nur77
NFAT:AP1
−log₂ P−value
log Enrichment
Y1
O1
IRF2
PU.1:IRF8
PRDM1
bZIP:IRF
IRF:BATF
Y2
O2
IRF1
bZIP:IRF
PRDM1
PU.1-IRF
IRF:bZIP
IRF:BATF
ATF3
Y3
O3
BATF
IRF4
PU.1-IRF
O
CTCF−SatelliteElement
GFX
OCT:OCT
ZNF317
EFL−1
Y4
O4
RUNX1
IRF4
Y5
O5
Y6
O6
SMAD4
SMAD2
ERG
RUNX1
Gfi1b
Tcf4
TCFL2
PGR
GATA
PQM−1
B
GO Biological Pathways (< 35 year old)
T cell activation
lymphocyte differentiation
response to interferon−gamma
response to ketone
leukocyte cell−cell adhesion
connective tissue development
ossification
eye development
MAP kinase activity
axonogenesis
leukocyte cell−cell adhesion
interleukin−4 production
Unique Y1 Y3 Y4 Y5 Y6 Y2
p.adjust
0.005
0.010
0.015
Gene Ratio
0.025
0.050
0.075
GO Biological Pathways (> 65 year old)
synapse assembly
SMAD protein signal transduction
innate immune response
response to mechanical stimulus
lymphocyte differentiation
cell morphogenesis
GTPase activity
axonogenesis
forebrain development
ossification
T cell activation
leukocyte cell−cell adhesion
Unique O1 O3 O4 O5 O6 O2
p.adjust
0.01
0.02
0.03
0.04
Gene Ratio
0.05
0.10
0.15
C
Peak at 1.5 h
NR4A1
FOS
FOSL1
Peak at 6 h
IRF1
IRF4
IRF9
Progressively rising
ATF5
BATF
PRDM1
SMAD family
SMAD3
SMAD4
SMAD7
RPKM
Time (hours)
0 1.5 6 24 48

**Figure S5, related to Figures 2D and E. Relationship between differential accessibility of regulatory regions and transcriptome**

**A.** Aggregate genome accessibility tracks of representative sites of cluster 7-10 shown in the heat plot in Figure 2D (left panel). Cyan-shaded areas indicate peaks that are more open in young (cyan) compared to older adults (magenta). Transcript data from corresponding genes as determined by RNA-seq (right panel). **B.** LogFC difference of transcripts differentially expressed between young and older adults from 48-hour RNA-seq data is plotted against logFC difference of differential peaks from ATAC-seq data annotated to the same gene at the indicated time point. Data are fitted with linear regression.

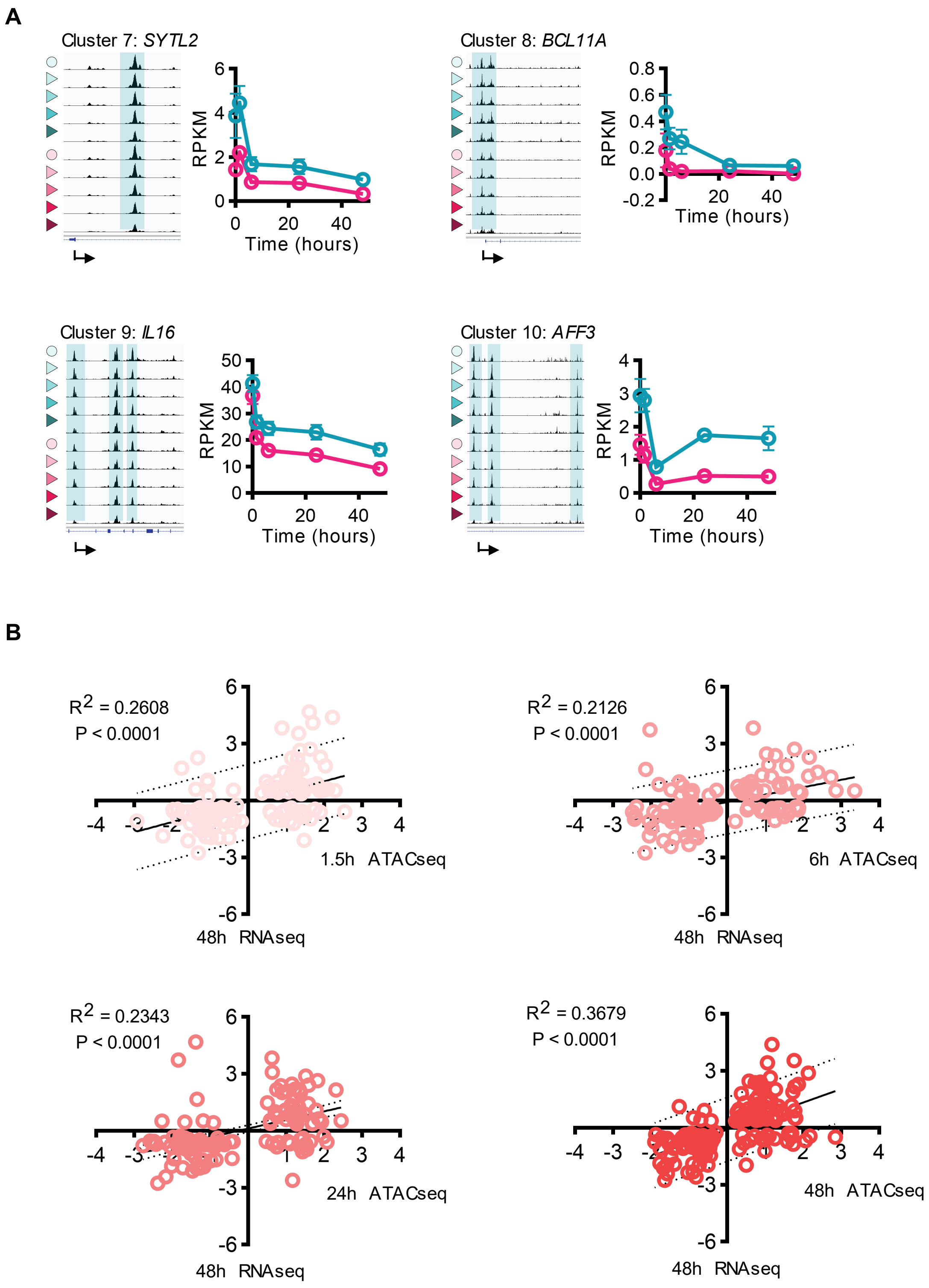

**Figure S6, related to Figure 4. Single cell epigenetic and transcriptional analysis of naïve CD4⁺ T cell responses.**

Nuclei pooled from unstimulated and activated naïve CD4⁺ T cells were subjected to scATAC- and scRNA-seq. Imputed *CD69* gene expression is projected on the UMAP of scATAC-seq. Clusters of resting and activated cells were distinguished based on containing *CD69*-expressing cells (left panel). UMAP maps of scATAC-seq and scRNA-seq data are shown for activated cells (right panels).

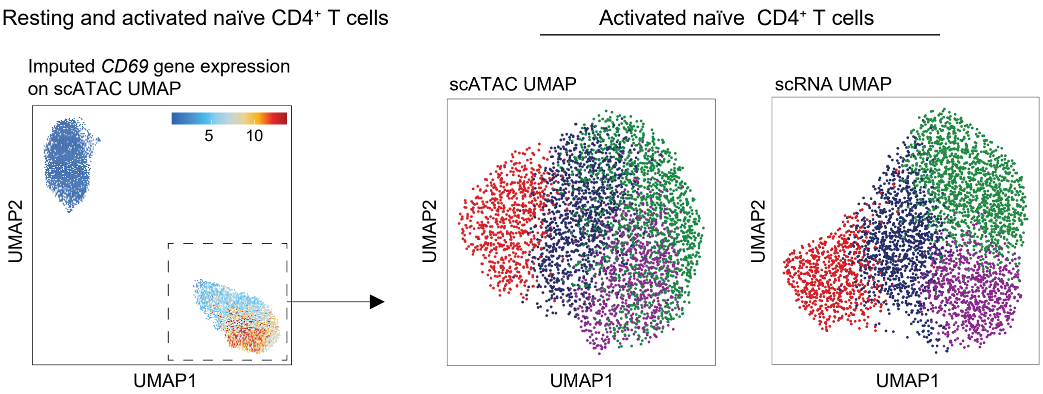

**Figure S7. Constitutive expression of the IL-2 receptor on naïve CD4+ T cells from older individuals.**

**A.** Gating strategy for naïve CD4+ T cells. **B.** *IL2RA* gene expression was quantified in unstimulated naïve CD4+ T cells from 10 young and 12 older adults. **C.** CD25 was stained on resting naïve CD4+ T cells from 12 young and 10 older adults. Data from B and C were analyzed with unpaired t-test. **p < 0.01.

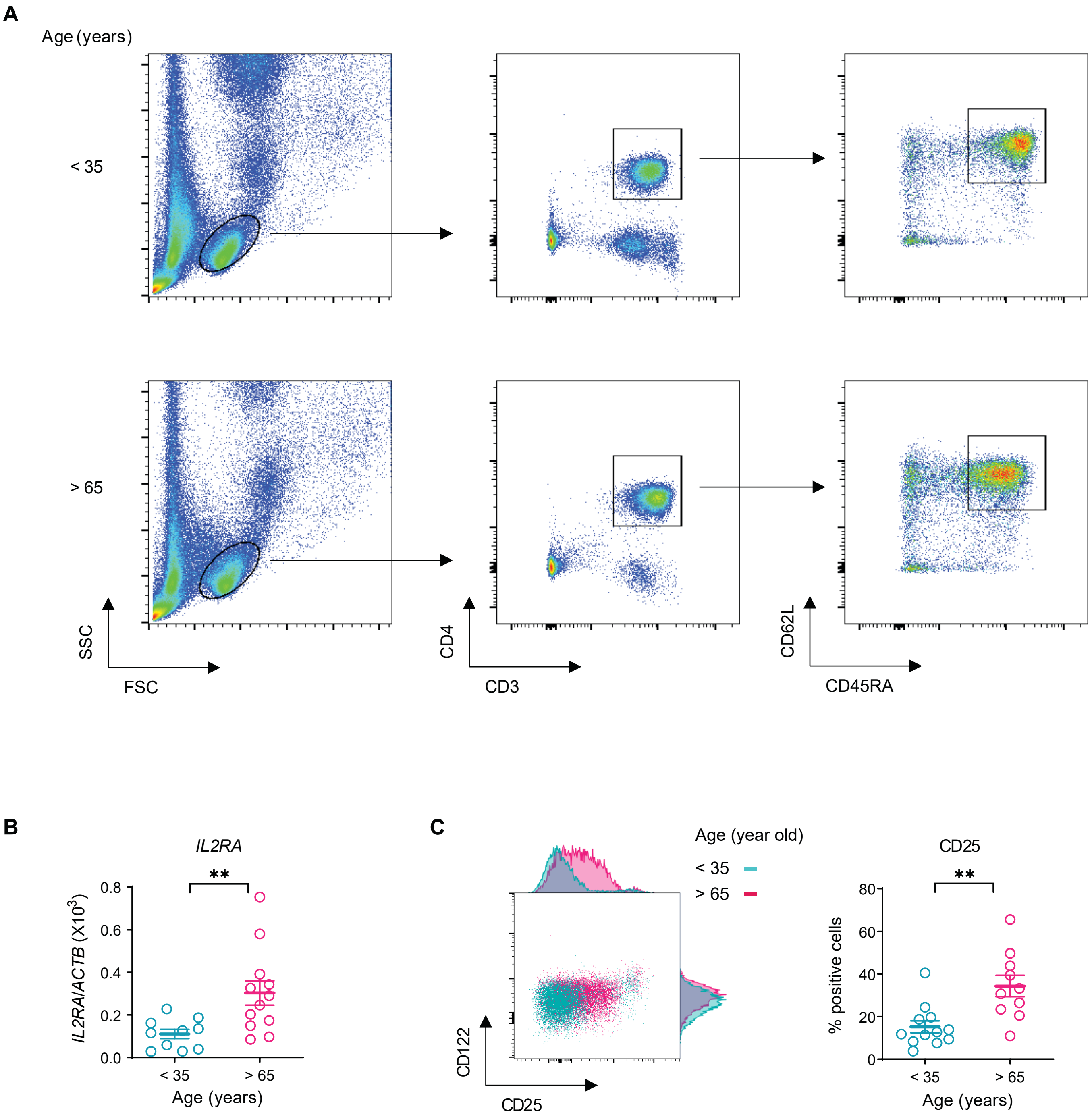

**Figure S8, related to Figure 5. IKZF2 binding to *IL2RA* enhancers from ENCODE ChIP-seq.**

**A.** ATAC-seq genome tracks of unstimulated naïve CD4$^{+}$ T cells from young and older adults were aligned with ENCODE ChIP-seq genome tracks of indicated TFs at *IL2RA* enhancer locus (see Figure 5A). The location of enhancer A and enhancer B is indicated by red bars. **B.** *IKZF2* and *IL2RA* transcripts from sorted CD25+ and CD25- naïve CD4$^{+}$ T cells from 7 donors.

**A**

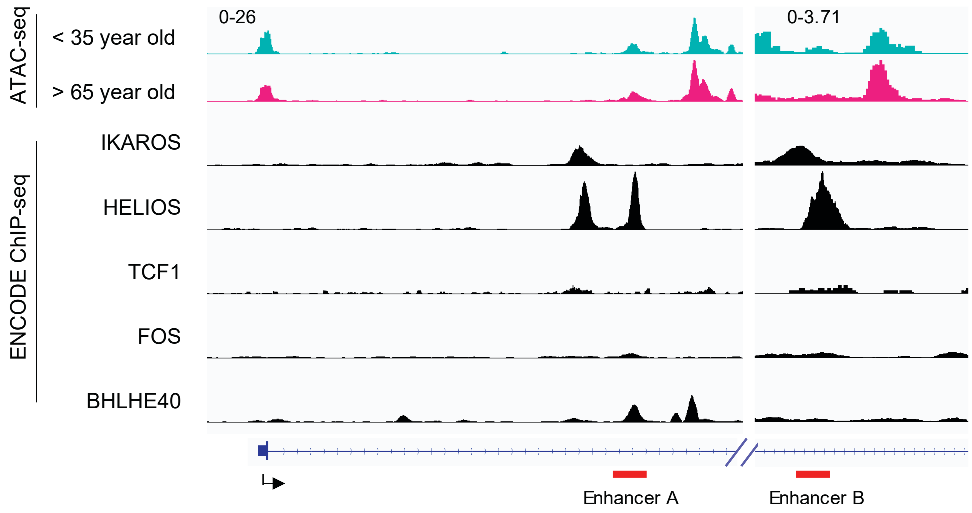


**B**

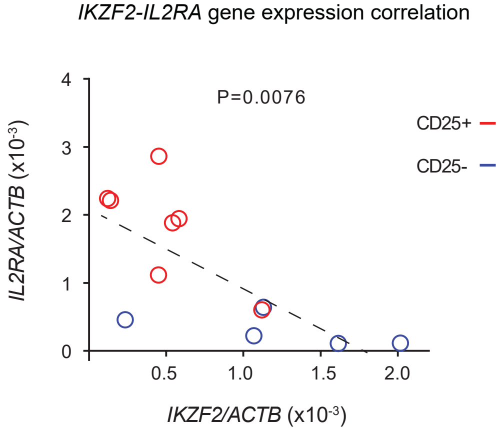

**Figure S9, related to Figure 6. Genome accessibility changes induced by IL-2 receptor blocking.**

**A.** LogFC of differential peaks between IL-2 receptor (IL-2R) blocking and PBS treatment (p<0.05) are plotted against average peak size. Peaks that are also differentially accessible between young and older individuals are indicated in dark red. **B.** Accessibility tracks of representative genes. Magenta shaded areas indicate peaks that are more open in older adults and with PBS treatment; cyan indicate peaks that are more open in young adults and with IL-2R blocking treatment.

A

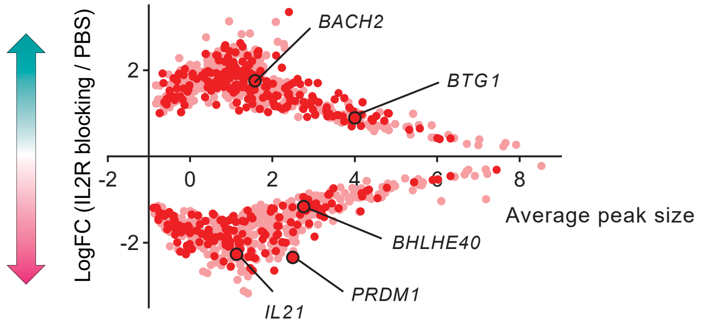


B

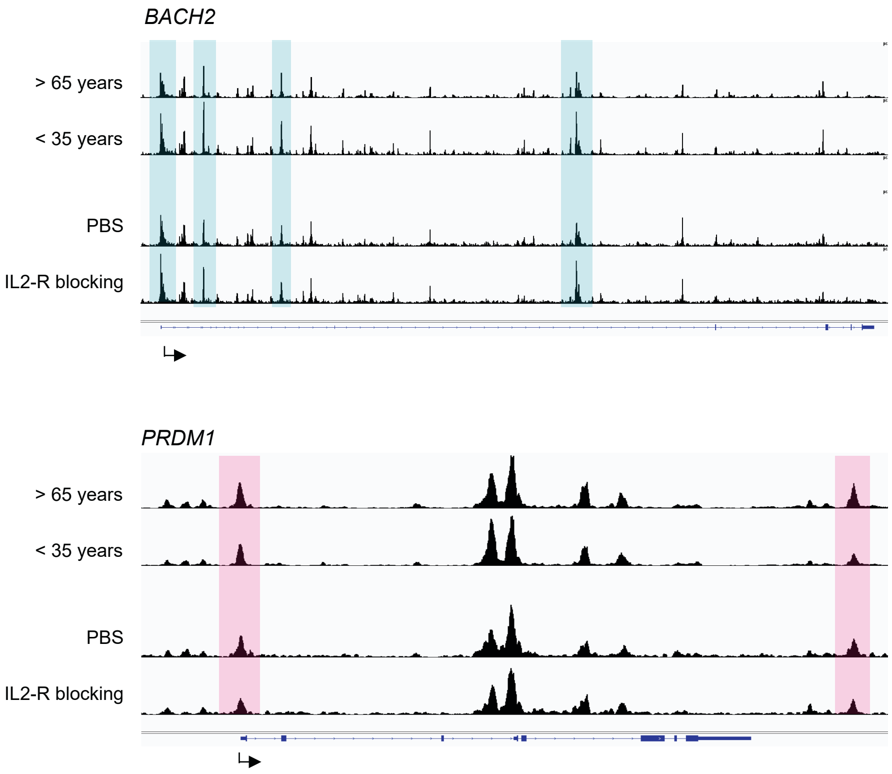

Supplement table 1. Study Population

| Subject | Ethnicity | Gender | Age | Enrolled experiment |
|---|---|---|---|---|
| 1 | Caucasian | M | 25 | TCR-stimulation time course |
| 2 | Asian | F | 70 | TCR-stimulation time course |
| 3 | Caucasian | M | 71 | TCR-stimulation time course |
| 4 | Caucasian | F | 30 | TCR-stimulation time course |
| 5 | Asian | F | 32 | TCR-stimulation time course |
| 6 | Caucasian | M | 65 | TCR-stimulation time course |
| 7 | Caucasian | M | 29 | TCR-stimulation time course |
| 8 | Caucasian | M | 74 | TCR-stimulation time course |
| 9 | Caucasian | F | 35 | scMultiome |
| 10 | Asian | M | 34 | scMultiome |
| 11 | Caucasian | F | 79 | scMultiome |
| 12 | Caucasian | M | 69 | scMultiome |
| 13 | Caucasian | M | 78 | STAT5 inhibition ATAC-seq |
| 14 | Caucasian | F | 80 | STAT5 inhibition ATAC-seq |
| 15 | Caucasian | M | 78 | STAT5 inhibition ATAC-seq |
| 16 | Caucasian | F | 81 | STAT5 inhibition ATAC-seq |

Supplement table 2. Antibodies

| Antibody name | Clone | Supplier | Catalog # | Application |
|---|---|---|---|---|
| Anti-CD69 BV605 | FN50 | BioLegend | 310937 | Surface staining |
| Anti-CD25 BV421 | BC96 | BioLegend | 302629 | Surface staining |
| Anti-CD3 PE-Cy7 | UCHT1 | BioLegend | 300420 | Surface staining |
| Anti-CD4 AF700 | RPA-T4 | BioLegend | 300526 | Surface staining |
| Anti-CD45RA AF750 | HI100 | BioLegend | 304151 | Surface staining |
| Anti-CCR7 PerCP-Cy5.5 | G043H7 | BioLegend | 353219 | Surface staining |
| Anti-ZAP70 pY319 AF647 | 17A/P-ZAP70 | BD | 557817 | Phosflow |
| Anti-SLP76 pY128 AF488 | J141-668.36.58 | BD | 558439 | Phosflow |
| Anti-ERK pT202 PE-CF594 | 20A | BD | 562644 | Phosflow |
| Anti-TCF1 PE | 7F11A10 | BioLegend | 65520725 | TF staining |
| Anti-BLIMP1 APC | 646702 | R&D systems | IC36081A | TF staining |
| Anti-IRF4 | polyclonal | Cell Signaling | 4964 | Western blot |
| Anti-BHLHE40 | Polyclonal | Novus Bio | NB100-1800 | Western blot |
| Anti-β-actin | 13E5 | Cell Signaling | 4970 | Western blot |
| Anti-ERK pT202 | D13.14.4E | Cell Signaling | 4370 | Western blot |
| Anti-LAT pY132 | polyclonal | Invitrogen | 44-224 | Western blot |
| Anti-STAT5 pY694 | polyclonal | Cell Signaling | 9351 | Western blot and ChIP |
| Anti-HELIOS | D8W4X | Cell Signaling | 42427 | Western blot and ChIP |
| Anti-CD25 | BC96 | BioLegend | 302602 | IL-2 receptor blocking |
| Anti-CD122 | TU27 | BioLegend | 339002 | IL-2 receptor blocking |
